# In vivo mitochondrial matrix proteome profiling reveals RTN4IP1/OPA10 as an antioxidant NADPH oxidoreductase

**DOI:** 10.1101/2021.10.14.464368

**Authors:** Isaac Park, Kwang-eun Kim, Jeesoo Kim, Subin Bae, Minkyo Jung, Jinhyuk Choi, Chulhwan Kwak, Myeong-Gyun Kang, Chang-Mo Yoo, Ji Young Mun, Kwang-Hyeon Liu, Jong-Seo Kim, Jae Myoung Suh, Hyun-Woo Rhee

## Abstract

Targeting proximity labeling enzymes to specific cellular locations is a viable strategy for profiling subcellular proteomes. Here, we generated transgenic mice expressing a mitochondrial matrix-targeted ascorbate peroxidase (MAX-Tg) to analyze tissue-specific matrix proteomes. Desthiobiotin-phenol labeling of muscle tissues from MAX-Tg mice allowed for efficient profiling of mitochondrial-localized proteins in these tissues. Comparative analysis of matrix proteomes from MAX-Tg muscle tissues revealed differential enrichment of mitochondrial proteins related to energy production in between different muscle groups. Reticulon 4 interacting protein 1 (RTN4IP1), also known as Optic Atrophy-10 (OPA10), was highly enriched in the cardiac and soleus muscles and was found to localize to the mitochondrial matrix via a strong mitochondrial targeting sequence at its N-terminus. Protein structure analysis revealed that RTN4IP1 is an NADPH oxidoreductase with structural homology to bacterial quinone oxidoreductase. Enzymatic activity assays, interactome analysis, and metabolite profiling confirmed a function for RTN4IP1 in coenzyme Q (CoQ) biosynthesis. *Rtn4ip1*-knockout C2C12 cells had reduced CoQ9 levels, were vulnerable to oxidative stress, and had decreased oxygen consumption rates and ATP production. Collectively, RTN4IP1 is a mitochondrial antioxidant NADPH oxidoreductase supporting oxidative phosphorylation activity in muscle tissue.

## Main

In multicellular organisms, each organ performs unique metabolic processes corresponding to a different proteome inventory ^1, 2^. As a central regulator of cellular metabolism, the mitochondrial proteome is expected to vary among tissues to meet tissue-specific metabolic demands. In support of this notion, imaging studies show significantly different mitochondrial ultrastructures among tissues ^3^, which are likely linked to distinct mitochondrial proteomes. In another example, the well-characterized oxidative phosphorylation (OXPHOS) complex, localized in the matrix and inner mitochondrial membrane (IMM) along with ATP synthase, is highly expressed in muscle tissues ^4^. However, the underlying details of the muscle-specific mitochondrial matrix proteome remain unclear. For example, a muscle-specific mitochondrial protein that scavenges elevated mitochondria reactive oxygen species (ROS) has not yet been identified ^5^.

Conventional approaches to identify the tissue-specific mitochondrial proteome are largely based on the subcellular fractionation and/or affinity purification followed by mass spectrometry ^6, 7 8, 9^. Previous studies using these conventional approaches showed differential expression levels of the mitochondrial proteome between tissues. Although these studies reported a tissue-specific enriched mitochondrial proteome in the heart, liver, kidney, and muscles, the mitochondrial proteome lists even within the same organ (e.g., muscle tissues) are discordant among studies, due to technical limitations such as the inevitable contamination of non-mitochondrial proteins using conventional purification methods ^10^. Furthermore, these studies cannot provide the sub-mitochondrial spatial resolution in proteome profiling. Although sub-organelle localization information can be revealed by conventional methods such as a protease K digestion assay for individual proteins, these methods suffer from technical artefacts introduced by harsh lysis conditions, which lead to unreliable and inconsistent findings ^11–13^. Consequently, accurate information regarding the mitochondrial proteome from different tissues at the sub-organelle level are lacking due to a lack of suitable methodology.

We reasoned that the engineered ascorbate peroxidase (APEX) technique can help overcome previous technical limitations in sub-organellar proteome analysis as APEX is capable of *in situ* biotinylation of the local proteome in which it is engineered to be expressed ^11, 14–16^. Since APEX-mediated proximity labeling results in the covalent biotinylation of tyrosine residues of endogenous proteins ^11, 14^, liquid chromatography and tandem mass spectrometry (LC-MS/MS) analysis of biotin-modified sites on endogenous proteins can reveal sub-cellular proteomes to which APEX is localized ^11^. Using proximity labeling techniques, we and our colleagues have successfully identified the sub-mitochondrial proteome of the mitochondrial matrix ^11, 14^, intermembrane space (IMS) ^17, 18^, and mitochondrial-associated membrane ^19, 20^ in the immortalized human cell line HEK293T, which is a commonly used *in vitro* model for mitochondrial proteome analysis using APEX2 or BioID.

Notably, the sub-mitochondrial proteome information using APEX in HEK293T cells^14, 17^, have been incorporated to recent updates of MitoCarta^21, 22^ which is a prominent resource curating 1136 human mitochondrial proteins along with their sub-mitochondrial localization information, pathway annotations and tissue-specific expression.^14, 17, 23^ However, it is questionable whether the sum of the data from different biological systems (e.g. HEK293T cells^14, 17^ and mouse tissues^23^) and methods (e.g. APEX^14, 17^ and fractionation^23^) in MitoCarta sufficiently reflects the diversity of *in vivo* physiological contexts such as tissue-specific sub-mitochondrial protein information. For example, creatine kinase B^24^ and TNAP^25^ are not found in the Mitocarta database although they are brown fat specific IMS proteins.^24, 25^ Therefore, new analytical methods that can provide direct experimental evidence for tissue-specific sub-mitochondrial proteomes are highly desirable.

Here, we extend the utility of APEX labeling for profiling sub-cellular proteomes to whole animal models. To this end, we establish a transgenic (Tg) mouse model that expresses mitochondrial <u>**m**</u>atrix-targeted engineered <u>**A**</u>PE<u>**X**</u>2 (MAX) to enable *in situ* biotinylation of the mitochondrial matrix-localized proteins of different tissues. Using this MAX-Tg mouse model, we demonstrate accurate and reproducible analysis of muscle-specific mitochondrial matrix proteomes and identify reticulon 4 interacting protein 1 (RTN4IP1) as a mitochondrial matrix-localized protein with antioxidant activity in muscle tissues.

## Results

### Generation of MAX-Tg mice for matrix-specific labeling in mouse tissues

To generate transgenic mice with constitutive expression of APEX2, a proximity labeling enzyme with higher specific activity than APEX ^16^, localized to the mitochondrial matrix by fusing the mitochondrial-targeting sequence (MTS) from COX4I1 (**Fig. 1a**). MAX-Tg mice were mated with wild-type (WT) mice and pups were born at the expected Mendelian ratio.

**Fig. 1:**
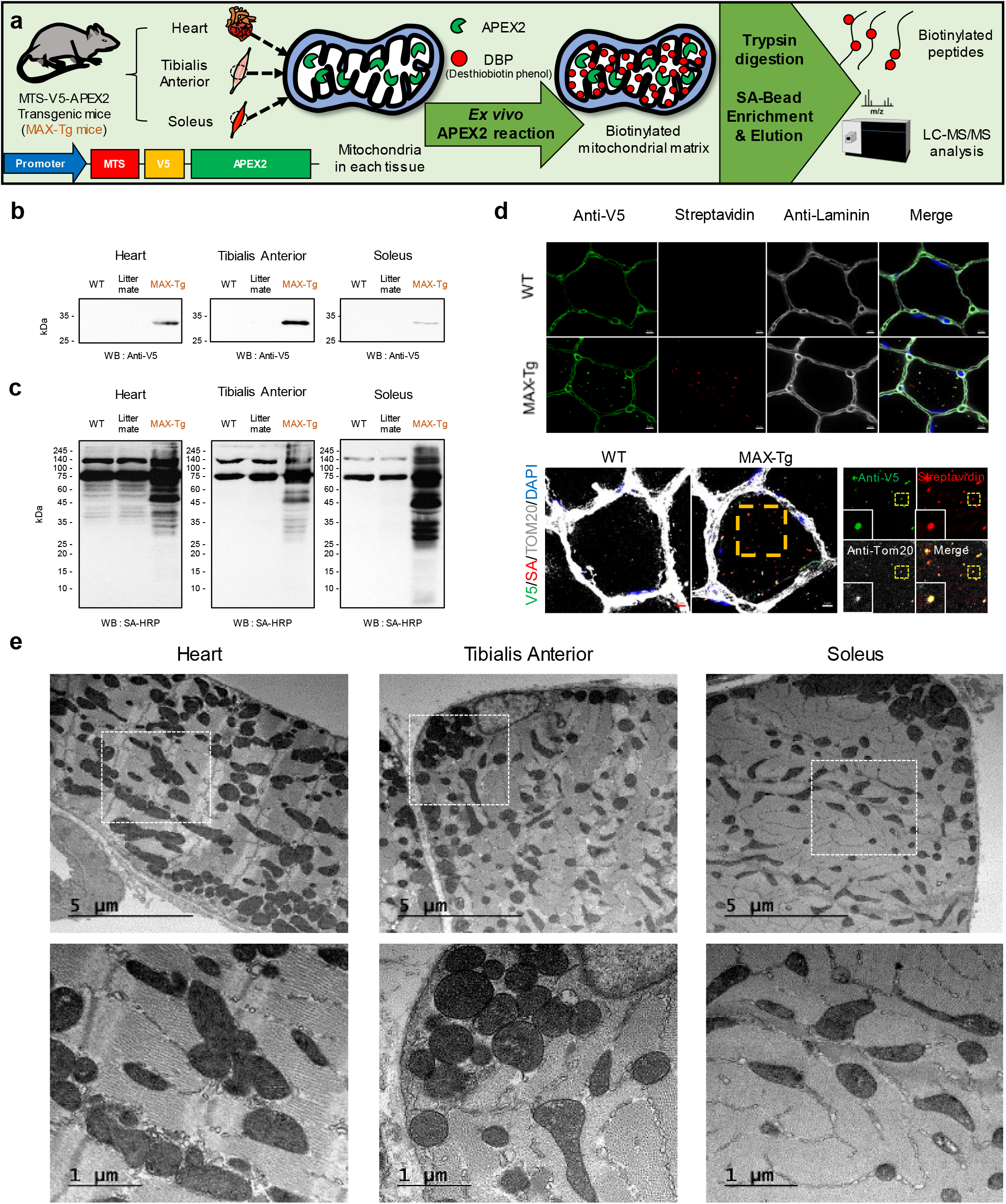
MTS-APEX2 transgenic (MAX-Tg) mice enable *in situ* profiling of mitochondrial matrix-specific proteomes. (a) Scheme for tissue-specific mitochondrial matrix proteome mapping using MAX-Tg mice. (b) Western blotting of MTS-V5-APEX2 (expected processed molecular weight: 28 kDa) in WT, littermate, and Tg mice. (c) Streptavidin (SA)-HRP western blotting of biotinylated proteins in WT, littermate, and Tg mouse tissues after APEX-mediated *in situ* biotinylation reaction (i.e., DBP and H_2_O_2_ treatment). (d) Confocal microscopy imaging of MTS-APEX2 *in situ* biotinylation in the TA muscle of MAX-Tg mice. Scale bar: 5 μm. (e) TEM of the mitochondrial matrix expression pattern of MTS-APEX2 in each muscle tissue of MAX-Tg mice. Scale bars: 5 μm (upper), 1 μm (lower).

Immunoblotting of desthiobiotin-phenol (DBP)-labeled muscle tissue lysates from MAX-Tg mice with anti-V5 and horseradish peroxidase (HRP)-conjugated streptavidin (SA; targeting biotinylated proteins) confirmed that MTS-V5-APEX2 was expressed and is functional in heart and skeletal muscle (**Fig. 1b,c**). Immunofluorescence imaging of heart and skeletal muscle tissues of MAX-Tg mice was performed to examine the subcellular localization of the APEX2-V5-APEX2 protein. While anti-V5 antibody showed non-specific binding to the surface of muscle myofibers in both WT and MAX-Tg mice, specific anti-V5 antibody staining within the muscle myofibers was observed only in MAX-Tg mice, which overlapped with that of the anti-TOM20 antibody staining of mitochondria and anti-biotin staining by streptavidin (**Fig. 1d**).

Transmission electron microscopy (TEM) was performed to assess whether MTS-V5-APEX2 specifically targets to the mitochondrial matrix compartment in MAX-Tg mice. APEX2 can be stained with 3,3′-diaminobenzidine (DAB) in fixed tissues used for TEM analysis ^26^ and DAB staining patterns from MAX-Tg muscle tissues confirmed specific targeting of APEX 2 to the mitochondrial matrix in MAX-Tg mice (**Fig. 1e**). The MTS-APEX2-stained aligned cristae structure was expressed in the heart and each skeletal muscle, and the structure of the fiber zone arranged around the mitochondria was in agreement with the morphology of mitochondria in healthy muscle ^27^. Mitochondria were mainly located between myofibrils, although some clustered together beneath the sarcolemma of muscle cells, which is consistent with the reported mitochondrial distribution in muscle ^27, 28^. These data indicate that MTS-V5-APEX2 is expressed in the MAX-Tg mouse muscle and displays proximity labeling activity in the mitochondrial matrix of muscle tissues.

### Divergent matrix proteome between mouse muscle tissues and a human cell line

Next, LC-MS/MS analysis was performed and muscle-specific mitochondrial matrix proteins were identified using the advanced Spot-ID technique ^11^ with *ex vivo* DBP labeling of 7-week-old Tg mouse tissues [heart, tibialis anterior (TA), and soleus muscles] (**Extended Data Fig. 1a**). To eliminate DBP-labeled peptides due to possible endogenous peroxidase activity in the muscles, the DBP-labeled mitochondrial proteome from each muscle tissue of MAX-Tg mice was compared with that from WT mice (negative control not expressing MTS-APEX2); results from triplicate analysis were quantitatively filtered according to P < 0.05 and fold change (FC) > 2 (*t*-test) (**Extended Data Fig. 1b**). Very low endogenous peroxidase activity was detected in the muscle tissues, with hundreds of DBP-labeled peptides identified by MTS-V5-APEX2 labeling in the muscle, heart, and cell line samples in a highly reproducible manner (**Supplementary Table 1**). Replicate samples from the same tissue origin (e.g., skeletal muscles, heart, HEK293T cells) showed high Pearson correlation coefficients in mass intensity and were well clustered (**Fig. 2a**). As expected, HEK293T cell samples showed low Pearson correlation coefficients with muscle tissues (r < 0.38). In contrast, the heart and skeletal muscle showed higher Pearson correlation coefficients (r > 0.5) with each other than with the HEK293T samples but formed a distinct cluster. Interestingly, the TA and soleus muscles, which are both skeletal muscles, also formed distinct clusters. These results indicate that the MTS-V5-APEX2-labeled proteome resolves the unique composition of the mitochondrial matrix proteome from different muscle groups.

**Fig. 2:**
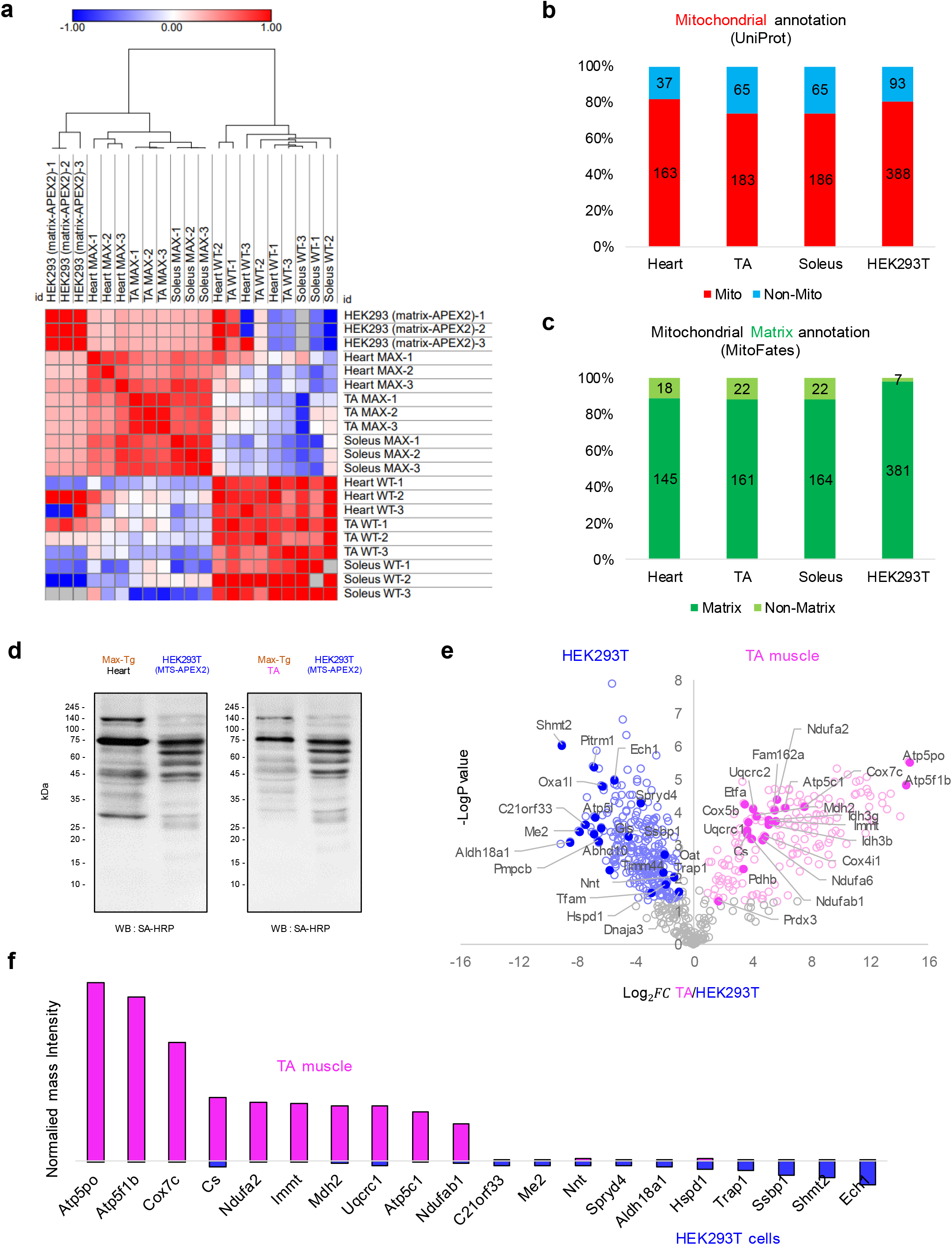
Mouse muscle tissues and human cell lines possess divergent mitochondrial matrix proteomes. (a) Heatmap of correlations between mass signal intensities of each replicate sample from WT mice, MAX-Tg mice, and HEK293T cells (heart, tibialis anterior, soleus, HEK293T cells stably expressing MTS-APEX2). Pearson correlation coefficients were calculated from each comparison. Hierarchical clustering was performed based on the Pearson correlation coefficients. See **Supplementary Table 2** for detailed information. (b) Mitochondrial distribution of the DBP-labeled proteins by MTS-APEX2 in each muscle tissues of MAX-Tg mice and HEK293T cells. Mitochondrial proteins were classified by annotation of their subcellular localization in UNIPROT. (c) Mitochondrial matrix distribution of the DBP-labeled proteins by MTS-APEX2 in indicated muscle tissues of MAX-Tg mice and HEK293T cells. Mitochondrial matrix proteins were classified with MitoFates prediction (see Methods). (d) Western blots of biotinylated proteins in MAX-Tg mouse tissues and HEK293T cells expressing MTS-V5-APEX2 after the APEX-mediated *in situ* biotinylation reaction (i.e., DBP and H_2_O_2_ treatment). (e) Volcano plot of the DBP-labeled proteome labeled by MTS-APEX2 in HEK293T cells (left) vs. TA muscle (right) from MAX-Tg mice. Statistical significance against fold change revealed significantly different proteins between the HEK293T proteome and TA muscle proteome. The top 20 DBP-labeled proteins based on the normalized mass intensities in each sample are marked with filled circles with their gene names. See **Supplementary Table 4** for detailed information. (f) Top 10 abundant DBP-labeled proteins labeled by MTS-APEX2 in TA tissue or HEK293T cells based on the normalized mass intensity.

In each filtered protein list, including 237 proteins in the TA muscle, 246 in the soleus muscle, and 193 in the heart, 71.9% of the proteins had a mitochondrial annotation in UniProt. We also employed MitoFates, a prediction method for cleavable N-terminal mitochondrial-targeting signals ^29^, to verify whether the filtered proteins were annotated as mitochondrial matrix proteins **(Fig. 2b,c**). Indeed, the MTS-V5-APEX2-labeled proteome was highly enriched (86.4%) for mitochondrial proteins as annotated by UniProt with predicted MTSs and no known IMS or outer mitochondrial matrix (OMM) protein was on the list except for RTN4IP1 (**Supplementary Table 3**). This indicated that MAX-Tg mice selectively label mitochondrial matrix proteins but not IMS or OMM proteins.

Cluster analysis of the mitochondrial proteome determined by MTS-APEX2 showed that the mitochondrial matrix proteomes of the TA muscle tissue and HEK293T cells ^11^ were the most distinct clusters (**Fig. 2a**). The biotinylated protein patterns observed by SA-HRP western blotting showed differences between the muscle tissues of MAX-Tg mice and cultured HEK293T cells (**Fig. 2d**). In the TA muscle, the most abundant proteins were related to functions in energy production, including OXPHOS complex proteins (e.g., NDUFAB1, NDUFA6, NDUFA2, UQCRC1, UQCRC2, COX4I1, COX7C, ATP5C1) along with several enzymes related to pyruvate regulation and the tricarboxylic acid (TCA) cycle (e.g., IDH3B, IDH3G, CS, MDH2) (**Fig. 2e,f**). Furthermore, mitochondrial calcium ion homeostasis-related proteins (LETM1, MCU) were significantly enriched in the TA muscle. Proteins related to mitochondrial RNA processing such as PTCD2 and PTCD3 were also relatively highly expressed in the muscle. Since PTCD2 and PTCD3 also play a role in the mitochondrial respiratory chain, this finding further reflects the dependence of TA muscle on energy production via respiration ^30, 31^.

In contrast, in HEK293T cells, mitochondrial chaperone proteins (e.g., DNAJA3, HSPD1) and proteins involved in transcription (e.g., SSBP1, TFAM) were strongly enriched by MTS-V5-APEX2 labeling (**Fig. 2e,f**). Several mitochondrial proteins involved in one-carbon metabolism (e.g., SHMT2) and glutamate metabolism (e.g., OAT, GLS, ALDH18A1), which play important roles in cancer metabolism ^32, 33^, were also highly enriched in the HEK293T cell mitochondrial matrix proteome. These data demonstrate that the composition of mitochondrial matrix proteomes in mouse muscle tissues are distinct from those in the immortalized HEK293T cell line.

### Different muscle tissues possess distinct matrix proteomes

Muscle fibers in different muscle groups have a distinct composition, which is expected to be reflected in their proteomes ^34^. We compared the MTS-APEX2-labeled proteome between the cardiac, TA, and soleus muscles. We filtered all muscle tissue MTS-APEX2-labeled proteomes according to MitoFates prediction ^29^ to focus on the mitochondrial matrix/IMM-localized proteome containing mitochondrial matrix-targeting sequences (see Methods). From the three muscle tissues, a total of 263 DBP-labeled proteins were filtered and representative proteins in each tissue are presented in **Fig. 3a**.

**Fig. 3:**
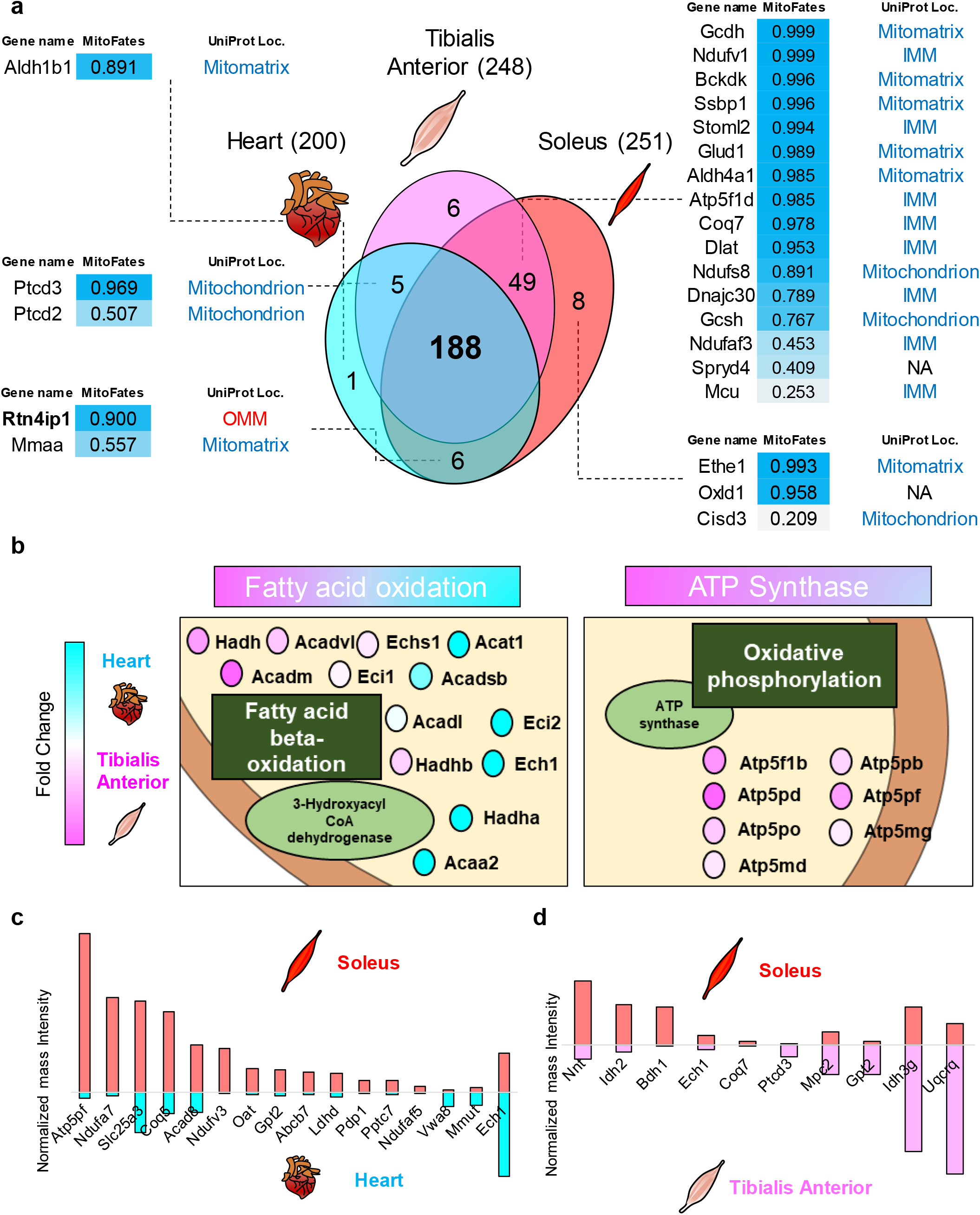
MAX-Tg mice resolve distinct matrix proteomes between muscle groups. (a) Venn diagram of identified mitochondrial matrix proteins from the heart, TA, and soleus tissues. Representative proteins are shown with the current subcellular informations in UNIPROT and MitoFates probability scores (see Methods and see **Supplementary Table 3** for detailed information). (b) Schematic representation of the tissue-enriched mitochondrial proteins with similar metabolic functions in the TA muscle and heart. Heart- and TA-enriched proteins are colored in light blue and pink, respectively. All proteins are color-coded to reflect the fold change of the normalized mass intensity of DBP-labeled peptides in the TA muscle and in the heart. Annotations with function and complex are based on UNIPROT and CORUM database. An expanded figure of tissue-enriched mitochondrial proteins (TA vs. heart) is shown in **Extended Data Fig. 2**. See **Supplementary Table 5** for detailed information. (c) Normalized mass intensities of asymmetrically DBP-labeled mitochondrial matrix proteins by MTS-APEX2 in the heart and soleus muscle. (d) Normalized mass intensities of asymmetrically DBP-labeled mitochondrial matrix proteins by MTS-APEX2 in the TA and soleus muscles.

Intriguingly, there were several tissue-specific mitochondrial matrix proteins, whereas 188 proteins were commonly identified across the three muscle tissues. Furthermore, quantitative comparative analysis between tissue proteomes revealed the differential enrichment of metabolic pathways in each tissue. We found higher expression levels of proteins related to the beta oxidation pathway (e.g., ACAA2, ACAT1, ACADSB, ECI2, ECH1, HADHA) in the heart compared to the TA muscle (**Fig. 3b** and **Extended Data Fig. 2**), which is consistent with the heart’s ability to produce energy from fatty acids; approximately 75% of the fatty acids are normally oxidized in mammalian hearts ^35^. Protein quality control system-related proteins (e.g., AFG3L2, PMPCA, CLPP) were also more abundantly expressed in the heart, whereas ubiquinone biosynthesis-related proteins such as ATP synthase complex and OXPHOS complex proteins were more highly expressed in the TA muscle (**Fig. 3b** and **Extended Data Fig. 2**). Similarly, a high relative abundance of ATP synthase complex and OXPHOS complex subunits was also found in the soleus skeletal muscle (**Fig. 3c**). Among proteins related to the TCA cycle, the expression levels of isotype proteins of isocitrate dehydrogenase (IDH) differed between the heart and muscle tissue: IDH3A, IDH3B, and IDH3G, which use NADH as a substrate ^36^, were abundant in the TA muscle, whereas IDH2, which uses NADPH ^36^, was more abundant in the heart (**Extended Data Fig. 2**).

The soleus muscle showed higher expression levels of mitochondrial proteins (i.e., NNT, IDH2) that utilize NADPH/NADP when compared with those of the TA muscle. The soleus muscle also showed higher expression levels of 5-demethoxyubiquinone hydroxylase (COQ7), which plays an essential role in coenzyme Q (CoQ) biosynthesis, whereas the TA muscle expressed higher levels of complex II and mitochondrial ribosome subunits (i.e., UQCRQ, PTCD3) and pyruvate-related proteins (i.e., GPT2 and MPC2**) (Fig. 3d)**. These data showed that each muscle tissue has a unique composition of mitochondrial matrix proteome.

### RTN4IP1 is a mitochondrial matrix protein

Unexpectedly, RTN4IP1 (also known as OPA10), which was previously reported as an OMM protein based on the conventional fractionation-based mitochondrial protein identification method ^37^, was identified as a matrix protein in the soleus and heart tissue samples labeled by MTS-APEX2. According to human genetic studies, RTN4IP1/OPA10 plays critical roles in mitochondrial function and is associated with optic neuropathy, muscle loss, and global developmental delay, which are related to mitochondrial dysfunction ^37, 38^. However, the molecular basis of RTN4IP1/OPA10 function in the mitochondria as well as its accurate sub-organellar localization are not clear. In this regard, we sought to clarify the sub-mitochondrial localization and identify the function of RTN4IP1/OPA10 in the mitochondria.

We first confirmed that the LC-MS/MS results obtained from MAX-Tg mice reflected the actual tissue proteome by evaluating RTN4IP1/OPA10 (hereafter RTN4IP1) expression in each tissue using antibody staining; RTN4IP1 expression was the highest in the heart followed by the soleus muscle, with the lowest expression in the TA muscle (**Fig. 4a**). We attempted solve the discrepancy in the sub-mitochondrial localization of RTN4IP1 between our result (matrix) and previous reports (OMM). MitoFates prediction with a high probability value (0.900) (**Fig. 3a**) as well as the unambiguous identification of biotinylated peptides in MTS-APEX2-labeled tissues strongly suggested its matrix localization. Immunofluorescence of RTN4IP1-V5-APEX2 in AD-293 cells also showed a clear mitochondrial expression patterns of both V5 and biotinylated proteins, which appeared to be plainly co-localized (**Fig. 4b**, top panel). We next tested the mitochondrial matrix-targeting capability of the MTS of RTN4IP1, predicted by MitoFates analysis, which is the N-terminal 32 amino acids; the peptide bond between R32 and D33 of the protein would be cleaved by mitochondrial processing peptidase. We expressed an APEX2 construct fused to the expected MTS (amino acids 1–32 of RTN4IP1) at the N-terminus of V5-APEX2 in AD-293 cells. As shown in the middle panel of **Fig. 4b**, confocal microscopy confirmed that both V5-stained and biotinylated proteins had a clear mitochondrial distribution pattern. We additionally established an MTS-deleted RTN4IP1-APEX2 construct [RTN4IP1(Δ1-32aa)-V5-APEX2], which showed a clear cytosolic distribution in both anti-V5 and SA staining (**Fig. 4b**, bottom panel). Collectively, the confocal imaging data strongly supported the matrix localization of RTN41P1 rather than the OMM.

**Fig. 4:**
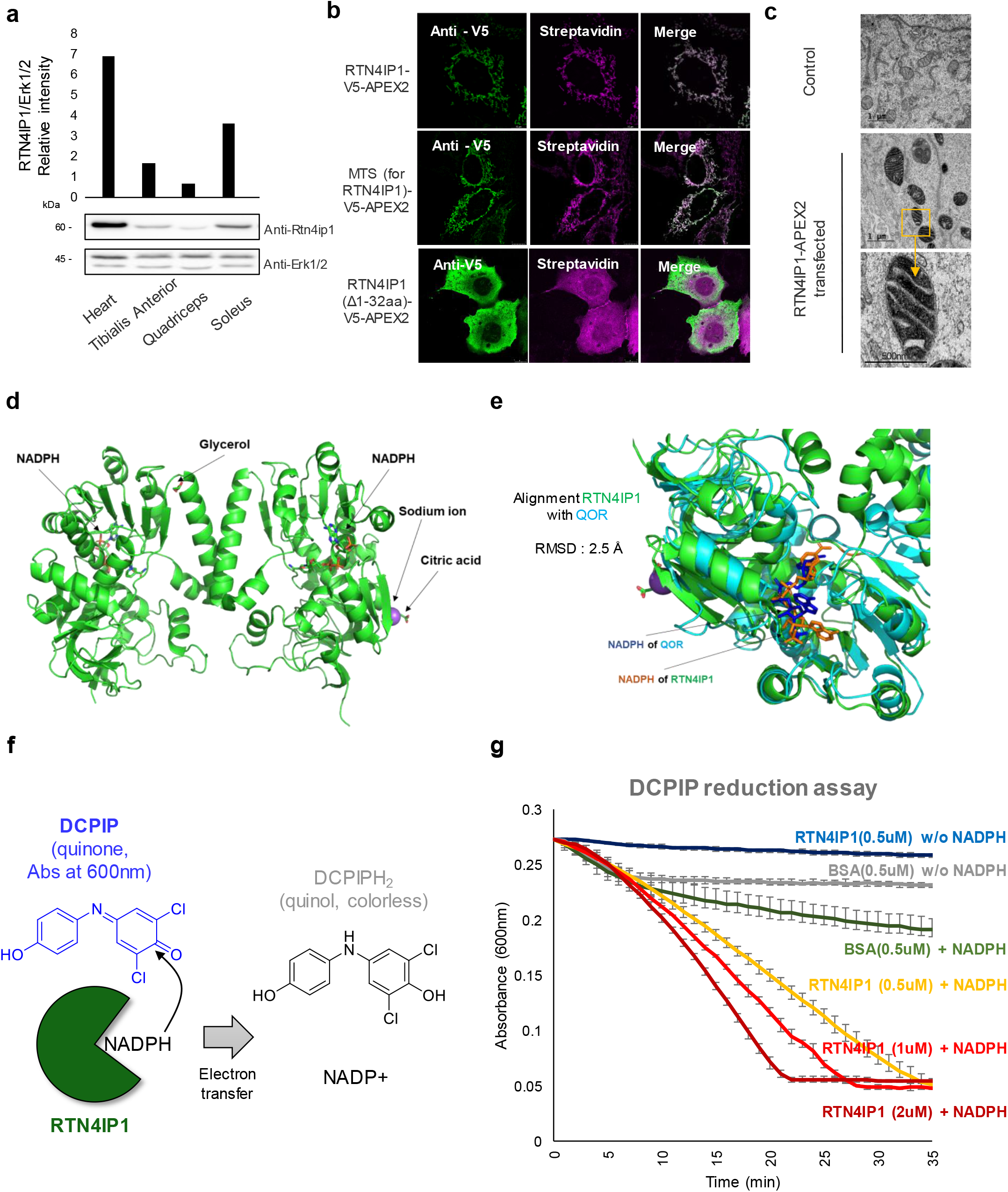
RTN4IP1/OPA10 targets the mitochondrial matrix and displays NADPH oxidoreductase activity. (a) Western blotting of the heart and three types of skeletal muscles. Anti-Erk1/2 was used as a loading control. The RTN4IP1/Erk relative intensity indicates the expression level of RTN4IP1 in various cell types and organs. (b) Confocal microscopy imaging of mitochondrial biotinylation by RTN4IP1-APEX2 in AD-293 cells. (pseudo-colored in Cy5) The N-terminal mitochondrial targeting sequence (MTS; ∼R32) of RTN4IP1 was predicted by MitoFates. (scale bar = 10 μm) (c) TEM of RTN4IP1-APEX2-transfected HEK293T cells (right) and untransfected HEK293T cells (left). (Scale bar: 2 μm) Both samples were treated with DAB and H_2_O_2_, followed by OsO_4_ staining. Mitochondrial matrix DAB/OsO_4_ staining of RTN4IP1-APEX2 is highlighted in the magnified images of the orange boxed region. (d) Crystal structure of RTN4IP1 (dimer form) from Protein Data Bank (PDB ID: 2VN8, green). (e) Comparison of the crystal structure between RTN4IP1 and quinone NADPH oxidoreductase (QOR) (PDB ID: 1QOR, blue). The molecular structure of co-crystalized NADPH is shown in the structure of RTN4IP1 and QOR. (f) Scheme for assay of the quinone oxidoreductase activity of RTN4IP1. (g) Real-time monitoring results using blue-colored oxidized dichlorophenolindophenol (DCPIP) as the quinone substrate (n=3). DCPIP turns colorless when it accepts an electron from NADPH.

Finally, we verified the sub-mitochondrial localization of RTN4IP1 by APEX-electron microscopy ^26^, which can unambiguously confirm the sub-mitochondrial localization of APEX-tagged proteins in contrast to conventional methods (i.e., protease restriction assay) ^10^. In RTN4IP1-APEX2-expressing cells, APEX-mediated DAB/OsO_4_ staining (darker regions) showed a staining pattern that is clearly consistent with localization to the mitochondrial matrix (**Fig. 4c**). Together, these results verified that RTN4IP1 is a mitochondrial matrix protein with a strong mitochondrial matrix-targeting sequence.

### RTN4IP1 is a mitochondrial NADPH oxidoreductase with antioxidant activity

RTN41P1 is an NADPH-binding protein according to Protein Data Bank (PDB) data; the co-crystal structure of RTN4IP1 with NADPH has been deposited in the PDB (ID: 2VN8) from a structural genomics study (**Fig. 4d**) without any report of its biochemical function. Considering that NADPH-binding proteins are rare in the mitochondrial matrix, and all known mitochondrial NADPH-binding proteins are related to biosynthetic pathways (i.e., IDH3G) or antioxidant control (i.e., NNT, TXNRD2), which are essential for supporting mitochondrial metabolism, we hypothesized that RTN4IP1 exerts its enzymatic activity by utilizing NADPH as a co-factor. In support of this notion, structural homology analysis for the reported structure of RTN4IP1 revealed high structural similarity (root mean square deviation 2.5Å for 1115 α-carbon) to *Escherichia coli* quinone NADPH oxidoreductase (QOR; PDB ID: 1QOR) ^39^ and that the two sequences share a conserved NADPH-binding quinone oxidoreductase domain ^40–42^ (**Fig. 4e**). This result indicated that RTN41P1 has a possible quinone oxidoreductase activity in the mitochondrial matrix.

To assess the NADPH oxidoreductase activity of RTN4IP1 with a quinone substrate, we purified recombinant RTN4IP1 protein and measured its catalytic activity using the oxidation/reduction indicator 2,6-dichloroindophenol (DCPIP); the absorbance spectrum (λ_max_ = 600 nm) of the oxidized “quinone” state of DCPIP is decreased when DCPIP is reduced to the “quinol” state by obtaining electrons from NADPH ^43–45^ (**Fig. 4f**). As shown in **Fig. 4g**, a rapid decrease in the absorption at 600 nm was observed after adding RTN4IP1 and NADPH to the sample compared with that of the control without RTN4IP1 or NADPH or the control with similar concentrations of bovine serum albumin and NADPH. These results indicated that RTN4IP1 plays a catalyst role in transferring electrons from NADPH to electron acceptors such as quinone molecules in the mitochondrial matrix.

### RTN4IP1 is required for CoQ biosynthesis

To further probe the detailed molecular function of RTN4IP1, we profiled the interactome of RTN41P1 in the mitochondrial matrix using TurboID ^46^, another widely used proximity labeling enzyme with a smaller labeling radius than APEX2. We introduced the RTN4IP1-TurboID and MTS-TurboID constructs to HEK293T-Rex cells, respectively. Proteins biotinylated by TurboID were profiled via definitive mass spectrometric identification of biotinylated peptides with the biotinylated lysine residues (K + 226 Da) ^47^. MTS-TurboID, which is evenly distributed in the mitochondrial matrix, was used as a control sample.

A total of 639 and 438 proteins were identified by MTS-TurboID and RTN4IP1-TurboID, respectively. Among these, 399 proteins were common between the two experiments (**Fig. 5a**). This large overlap is reasonable considering that both TurboID systems are localized in the mitochondrial matrix and the biotinylation of TurboID in the matrix space occurs in a somewhat promiscuous manner. To further narrow down the RTN41P1 interactome in the matrix, we performed quantitative comparative analysis between RTN41P1- and MTS-TurboID based on the normalized mass intensity (**Fig. 5b)**. From this analysis, we obtained 32 interactome candidate proteins that were reproducibly labeled by RTN4IP1-TurboID (P < 0.05, FC > 2; **Fig. 5b, Extended Data Fig. 3**). Among them, two proteins (COQ3, COQ5) are involved in CoQ biosynthetic processes; COQ3 was the most strongly biotinylated protein by RTN4IP1-TurboID, suggesting a probable role for RTN4IP1 in CoQ production pathway within the mitochondria (**Fig. 5c**).

**Fig. 5:**
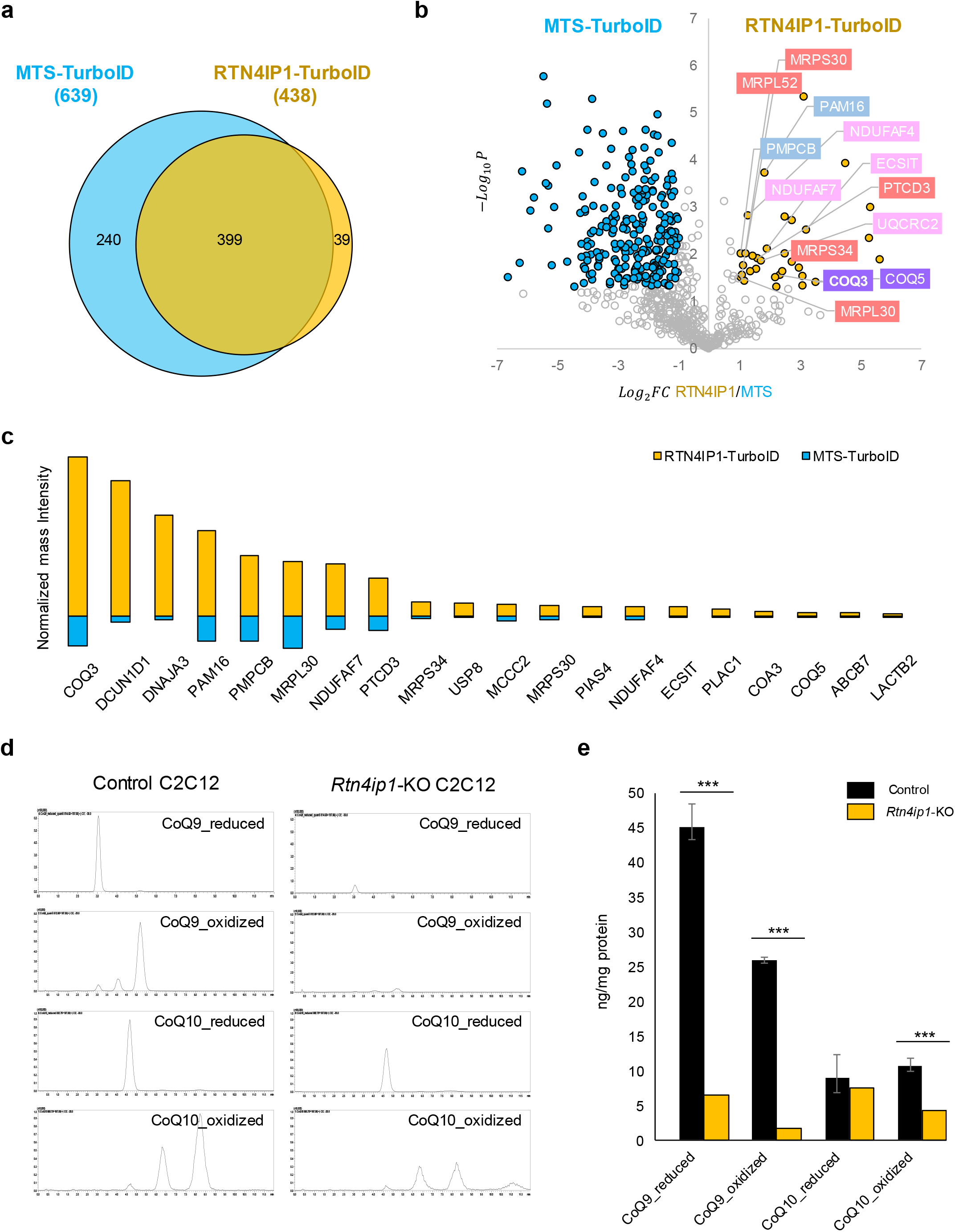
RTN4IP1/OPA10 is required for CoQ synthesis. (a) Number of identified biotinylated proteins by MTS-TurboID or RTN4IP1-TurboID in HEK293T cells via the Spot-BioID workflow. See **Supplementary Table 6** for detailed information. (b) Volcano plot of MTS-TurboID (left) vs. RTN4IP1-TurboID (right) biotinylated proteins. Statistical significance against fold change revealed significantly different proteins between MTS-TurboID-labeled and RTN4IP1-TurboID-labeled samples. A total of 32 proteins were significantly biotinylated by RTN4IP1-TurboID (RTN4IP1 interactome); P < 0.05, FC > 2. The proteins clustered by each function are highlighted. See **Extended Data Fig. 3** for details. (c) Normalized mass signal intensities of the top 20 biotinylated proteins labeled by RTN4IP1-TurboID among the 32 significantly biotinylated proteins. (d) LC-MS/MS analysis of coenzyme Q9 (CoQ9) in *Rtn4ip1*-knockout (KO) C2C12 cells (300 L/min flow rate; retention time for coenzyme Q9, 5.3 min; retention time for coenzyme Q9_reduced, 3.2 min). CoQ10 measurement using LC-MS (300 L/min flow rate; retention time for CoQ10, 8.5 min). (e) CoQ9 and CoQ10 levels in control and *Rtn4ip1*-KO C2C12 cells (n=6). *P < 0.05; **P ≤ 0.01, ***P < 0.001 (unpaired *t*-test).

Since CoQ and its intermediates include quinone moieties, and several steps of CoQ biosynthesis are believed to be assisted by electron transfer activity from as-yet unknown proteins in the matrix space ^48^, we hypothesized that RTN4IP1 is likely involved in the biosynthetic process of CoQ based on its localization (mitochondrial matrix), *in vitro* enzymatic activity (electron transfer from NADPH to quinone), and its interactome information in the mitochondrial matrix (COQ3, COQ5). To evaluate this possibility, we performed metabolite analysis in control and *Rtn4ip1*-knockout (KO) C2C12 mouse myoblast cell lines established using CRISPR/Cas9 genome editing (see Methods). Using targeted LC-MS analysis, we measured CoQ levels in control and *Rtn4ip1*-KO C2C12 cells and found a marked reduction in both the oxidized and reduced forms of CoQ9, the predominant form of CoQ in mice ^49^, in *Rtn4ip1*-KO cells compared to control (**Fig. 5d,e**). These results indicate an essential role for mitochondrial matrix protein RTN4IP1 in the biosynthetic pathway of CoQ.

CoQs mediate electron transport between Complex I/II and Complex III of the OXPHOS complex and also functions as an antioxidant ^50^; both functions of CoQ are beneficial for maintaining normal OXPHOS complex activity. Notably, CoQ metabolites were shown to have protective roles against oxidative stress ^50^. Thus, we conducted several experiments to determine whether the function of RTN4IP1 is related to antioxidant functions and regulation of OXPHOS complex activity in the mitochondria of muscle cells.

### RTN4IP1 deficiency induces oxidative stress

In the mitochondrial matrix, CoQ and tocopherol (vitamin E) have a quinone moiety, and both are well-characterized antioxidants in their reduced quinol state ^51^. Mitochondrial-synthesized CoQ can protect against excessive ROS generation in various subcellular membranes through the lipid transport machinery ^50^. Thus, we hypothesized that RTN4IP1 may protect the mitochondria from ROS by supporting the generation of protective CoQ using NADPH as a co-factor in the mitochondrial matrix.

We conducted TEM imaging to evaluate ultrastructural changes of subcellular components due to RTN4IP1 depletion. As shown in **Fig. 6a**, many vacuole-like structures were highly increased in *Rtn4ip1*-KO C2C12 cells. In particular, we observed striking structural aberrations of mitochondria in *Rtn4ip1*-KO cells. Mitochondrial cristae of *Rtn4ip1*-KO cells were collapsed, and a reduced electron density was observed in the matrix (**Fig. 6b**). Outer membrane rupture was observed in numerous mitochondria in *Rtn4ip1*-KO cells (**Fig. 6b, Extended Data Fig. 4**), with structures very similar to those observed under ferroptosis-inducing conditions ^52–54^. Since CoQ plays a protective role against ferroptosis ^50^, our data suggest that RTN4IP1 might be a potent suppressor of ferroptosis by maintenance of CoQ levels. In addition, a large number of multilamellar body structures, which are related to the autophagic process to degrade damaged mitochondria ^55^, were observed in *Rtn4ip1*-KO cells (**Fig. 6a, Extended Data Fig. 4a,b**).

**Fig. 6:**
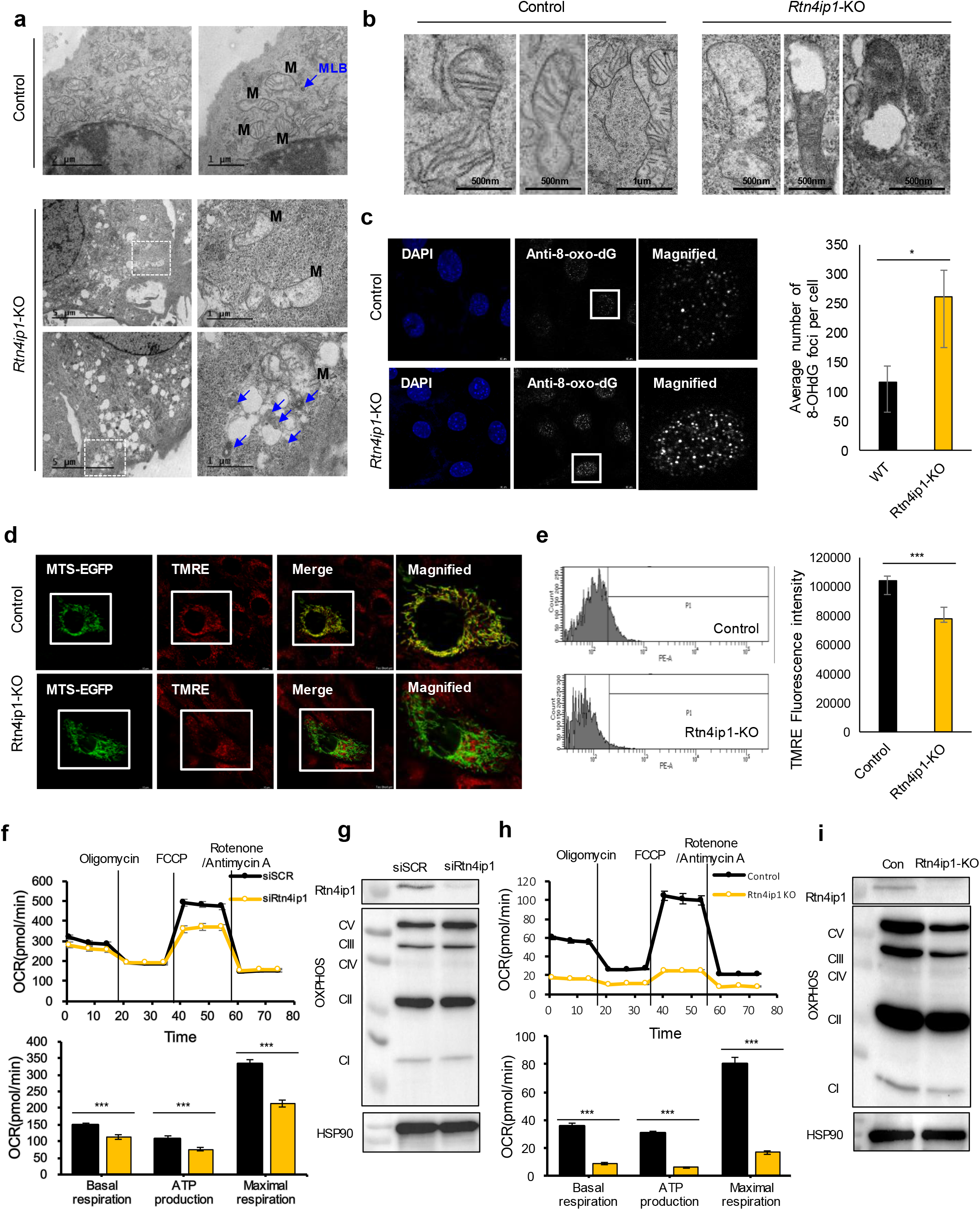
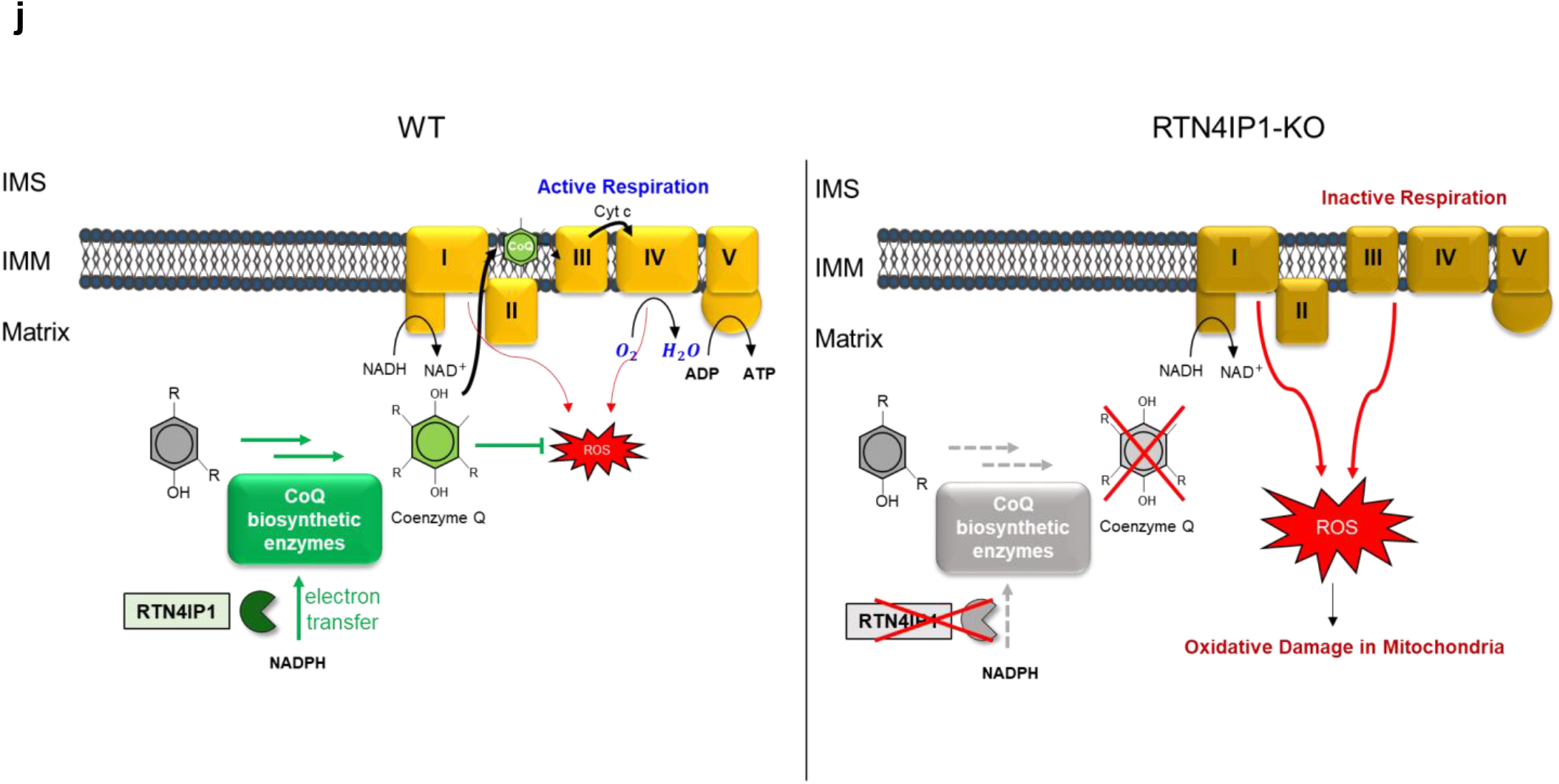
RTN4IP1/OPA10 regulates oxidative stress and mitochondrial respiration. (a) TEM of control (upper panel) and *Rtn4ip1*-KO (lower panel) C2C12 cells. Mitochondrial structures are marked with M and multilamella body (MLB) structure are marked with blue arrows. (b) Magnified TEM images of control (left panel) and *Rtn4ip1*-KO (right panel) C2C12 cells focusing on the mitochondria. (c) Confocal microscopy images of control and *Rtn4ip1*-KO C2C12 cells (scale bar = 10 μm). DNA regions oxidized by intracellular reactive oxygen species (ROS) were stained with an anti-8-oxo-dG monoclonal antibody and overall DNA was stained with DAPI. The average number of 8-OHdG foci per cell was counted in images from control and *Rtn4ip1*-KO C2C12 cells (n=10). *P < 0.05; **P ≤ 0.01, ***P < 0.001 (unpaired *t*-test). (d,e) Mitochondrial membrane potential measurement of control and *Rtn4ip1*-KO C2C12 cells by TMRE fluorescence and flow cytometry. (D) Confocal microscope imaging of TMRE and MTS-EGFP (a mitochondrial marker protein) in control and *Rtn4ip1*-KO C2C12 cells (scale bar = 10 μm). (E) Representative histograms of control and *Rtn4ip1*-KO C2C12 cells incubated with 1uM TMRE (left panel). 10,000 cells assayed for TMRE fluorescence by flow cytometry. Mean fluorescence of the C2C12 cells in each sample measured by microplate reader (right panel) (n=4). *P < 0.05; **P ≤ 0.01, ***P < 0.001 (unpaired *t*-test). (f) Measurement of oxygen consumption rate (OCR), basal respiration, maximal respiration, and ATP production in control (siSCR) and *Rtn4ip1* knockdown C2C12 cells (siRtn4ip1) (n=3). *P < 0.05; **P ≤ 0.01, ***P < 0.001 (unpaired *t*-test). (g) Western blotting with anti-RTN4IP1 and an antibody cocktail for the OXPHOS complex in siSCR and si*Rtn4ip1* C2C12 cells. (h) Measurement of OCR, basal respiration, maximal respiration, and ATP production in control and *Rtn4ip1*-KO C2C12 cells (n=3). *P < 0.05; **P ≤ 0.01, ***P < 0.001 (unpaired *t*-test). (i) Western blotting with anti-RTN4IP1 and an antibody cocktail for the OXPHOS complex in control and *Rtn4ip1*-KO C2C12 cells. (j) Proposed role of RTN4IP1 in CoQ biosynthesis and related functions.

Next, we assessed oxidative stress based on the oxidized nuclear DNA content in *Rtn4ip1*-KO cells using anti-8-oxo-dG monoclonal antibody (**Fig. 6c**). In control C2C12 cells, DNA staining with the antibody showed a lower number of foci (average 117 per cell) than that in *Rtn4ip1*-KO cells, with a larger number of foci (average 262 per cell) observed in the nucleus (**Fig. 6c**). Collectively, these results indicate that a RTN4IP1 deficiency leads to endogenous ROS accumulation and increased damage due to external oxidative stress, which may be attributed to a reduced levels of the antioxidant CoQ9.

### *Rtn4ip1*-KO cells show decreased membrane potential and OXPHOS activity

We further tested whether RTN4IP1 function is associated with OXPHOS activity of the mitochondria given the essential role of CoQ in the electron transport chain. We measured mitochondrial membrane potential using the fluorescent signal of tetramethylrhodamine, ethyl ester (TMRE), and confirmed that the TMRE signal of *Rtn4ip1*-KO cells was largely decreased compared to the detected level in control C2C12 cells (**Fig. 6d,e**).

As decreased membrane potential is directly associated with decreased OXPHOS activity in the mitochondria due to oxidative damage of electron transport chain proteins, we next evaluated the oxygen consumption rate (OCR) and expression of OXPHOS complex subunits. The OCR was reduced in *Rtn4ip1*-knockdown (KD) C2C12 cells transfected with small interfering RNA (siRNA) compared with that of control cells (**Fig. 6f**). Basal respiration, ATP production, and maximal respiration all decreased in *Rtn4ip1*-KD cells. This indicated that *Rtn4ip1* KD causes an abnormality in the respiratory system, further supporting the direct or indirect role of *Rtn4ip1* in the cellular respiration process. However, there were no differences in the levels of OXPHOS complex proteins between the *Rtn4ip1*-KD and control cells (**Fig. 6g**).

In *Rtn4ip1*-KO cells, the reduction in OCR was much more severe than that observed in *Rtn4ip1*-KD cells, and the expression levels of OXPHOS complex proteins, especially those in Complexes I, III, and V, decreased in *Rtn4ip1*-KO cells (**Fig. 6h, i**). Collectively, these results support that a deficiency in RTN4IP1 may reduce OXPHOS efficiency due to low levels of CoQ9 in muscle cells. Overall, we can propose that RTN4IP1 is involved in the biosynthesis of CoQ in the mitochondrial matrix, which is crucial for protection against ROS and the OXPHOS complex activity in the mitochondria (**Fig. 6j**).

## Discussion

We established MAX-Tg mice that constitutively express mitochondrial matrix-targeted APEX that enables profiling of tissue-specific for matrix proteomes. Tissue imaging and western blot analysis showed that MTS-APEX2 was efficiently expressed in the mitochondrial matrix of mouse muscle tissues with considerable peroxidase enzyme activity (based on DAB staining and *in situ* biotinylation). The robust APEX2 activity in *in vivo* muscle tissues is consistent with a recent report using APEX in the mouse brain for electron microscope imaging ^56^.

Using MAX-Tg for the analysis of muscle tissue mitochondrial matrix analysis, we observed that RTN4IP1 was highly enriched in the soleus/heart muscle tissue. RTN4IP1 was first proposed as a mitochondrial protein interacting with RTN4 (also known as NOGO) using a yeast two-hybrid assay ^57^. This interaction was confirmed by a co-immunoprecipitation experiment with the endoplasmic reticulum membrane protein RTN4, suggesting that RTN4IP1 localized at the OMM ^37^. However, the detailed molecular mechanisms of RTN4IP1 underlying its role in OXPHOS activity at the mitochondrial matrix/IMM ^37, 38^ are yet to be investigated. In the present study, we revealed that RTN4IP1 is mainly expressed in the mitochondrial matrix of the heart and soleus muscle tissues, and is involved in CoQ biosynthesis, which can explain its direct regulation of OXPHOS complex activity.

Interestingly, RTN4IP1-deficient cells were vulnerable to oxidative stress and they showed a similar mitochondrial morphology to that found under ferroptosis, which is a distinct form of cell death characterized by lipid peroxidation ^58, 59^. Notably, CoQ are well-characterized antioxidants to inhibit lipid peroxidation ^51^ and recent studies identified that ferroptosis suppressor protein 1 (FSP1), a NAD(P)H-dependent oxidoreductase plays a role to halt the propagation of lipid peroxides by production of reduced CoQ at the plasma membrane ^50, 60^. Since RTN4IP1 is also an NADPH oxidoreductase involved in the biosynthesis of CoQ, it can be considered as another major CoQ-dependent suppressor of ferroptosis. Experiments are currently underway to confirm this hypothesis. Our results may also provide insights into the treatment of patients with genetic defects of *RTN4IP1*. These patients suffer from optic neuropathy and ataxia ^37, 38, 61–63^, and our results suggest that delivery of antioxidant molecules or CoQ supplements may help to relieve these symptoms.

Notably, RTN4IP1 is a highly conserved protein from humans to other organisms. Rad-8—an ortholog of RTN4IP1 in *Caenorhabditis elegans*—was identified as a mitochondrial protein, with mutations of this gene causing increased sensitivity to oxidative stress and mitochondrial defects ^64^. The yeast ortholog of RTN4IP1, Yim1p, is also a mitochondrial NADPH quinone oxidoreductase protein (see https://www.yeastgenome.org/) and its mutant showed reduced resistance to the oxidative stress ^65^. Thus, it would be interesting to confirm whether these RTN4IP1 orthologs in other eukaryotic systems exhibit similar NADPH activity in CoQ biosynthesis. The similarity of predicted three dimensional structures of RTN4IP1 orthologs by Alphafold^66^ further support this idea (**Extended Data Fig. 5)**. In the International Mouse Phenotype Consortium dataset, whole-body knockout of RTN4IP1 (*Rtn4ip1*^*-/-*^) is lethal (see https://www.mousephenotype.org/), indicating an essential physiological role for RTN4IP1.

In summary, we successfully applied the APEX2 technique to generate MAX-Tg mice which can be used for obtaining tissue-specific mitochondrial matrix proteome data. Using these MAX-Tg mice, we identified new mitochondrial matrix proteins such as RTN4IP1, which is highly expressed in the soleus and cardiac muscles. Further experiments revealed that RTN4IP1 is an NADPH oxidoreductase that functions in antioxidant defense to support mitochondrial respiration activity. Overall, this study demonstrates that MAX-Tg mice are potentially useful for identifying tissue-specific mitochondrial proteins and their metabolism in diverse disease models.

## Methods

### Ethical statement

Animal studies were approved by the Institutional Animal Care and Use Committee of Seoul National University (no SNU180521-2-1). Mice were maintained in a specific pathogen-free institute.

### Matrix-V5-APEX2 Tg-MAX mouse model

#### Tg DNA sample preparation

DNA for microinjection of the Mito_V5_APEX2TG vector was prepared as follows. Mito_V5_APEX2 DNA is a fragment (∼2.1 kb) containing the cytomegalovirus (CMV) promoter with MluI and NaeI enzymes located at the front and back end of the promoter sequence for restriction enzyme digestion to secure the fragment. A backbone fragment of ∼2.6 kb and ∼1.3 kb was deleted through gel extraction, and a ∼2.1-kb fragment was used.

#### Generation of Tg mice

MTS-V5-APEX2 Tg mice were generated, interbred, and maintained in pathogen-free conditions at Macrogen, Inc. (Seoul, Republic of Korea). All manipulations were conducted under Macrogen Institutional Animal Care and Use Committee approval. Briefly, 5–8-week-old female C57BL/6N mice were intraperitoneally injected with pregnant mare serum gonadotropin (7.5 IU) and human chorionic gonadotropin (hCG; 5 IU) to induce superovulation. After hCG injection, the female mice were mated with C57BL/6N stud male mice. The next day, the presence of the virginal plug was checked in the female mice; those lacking the plug were sacrificed and the fertilized embryos were harvested. MTS-V5-APEX2 DNA was then co-microinjected into a single-cell embryo. Standard microinjection procedures were used for Tg mouse production (Macrogen, Inc.). DNA (4 ng/μL) for microinjection was directly injected into the male pronucleus of zygotes using a micromanipulator, and microinjected embryos were incubated at 37°C for 1–2 h. Fourteen to sixteen injected single-cell-stage embryos were surgically transplanted into the oviducts of pseudopregnant recipient ICR mice. After the F0 offspring were born, genotyping was performed using cut tail samples to detect the presence of the transgene, followed by polymerase chain reaction (PCR) confirmation of extracted genomic DNA with a specific primer pair (F: 5′-GTCGACGAGCTCGTTTAGTGA, R: AAGACCGTTGTTAGCGCTGTG-3′).

Mice were maintained under a 12 h light-dark cycle in a climate-controlled specific pathogen-free facility in Seoul National University. Mice were housed in same-sex groups and littermates of the same sex were randomly assigned to experimental groups. Standard chow diet and water were provided *ad libitum*.

### Cell culture and transfection

HEK293T and C2C12 cells were obtained from the American Type Culture Collection (Manassas, VA, USA), which were used at >20 passages. HEK293T Flp-in T-rex cells were obtained from Thermo Fisher Scientific (Cat. no. R78007). The cell lines were frequently checked and tested for morphology and mycoplasma contamination under a microscope but were not authenticated. All cell lines were maintained in high-glucose Dulbecco’s modified Eagle medium (DMEM) with 10% fetal bovine serum (FBS) at 37°C in 5% CO_2_ (v/v). All cell lines were transiently transfected at 60–80% confluence using polyethyleneimine (PEI).

### Expression plasmids

Genes were cloned into the specified vectors using standard enzymatic restriction digestion, and then ligated with T4 DNA ligase. To generate constructs in which short tags (e.g., V5 epitope tag) or signal sequences were appended to the protein, the tag was included in the primers used to PCR amplify the gene. The PCR products were then digested with restriction enzymes and ligated into cut vectors (e.g., pcDNA3, pcDNA5, and pET-21a). In all cases, the CMV promoter was used for expression in mammalian cells. The Extended data lists the genetic constructs cloned and used in this study.

### Histology and immunofluorescence staining

Mice were anesthetized with isoflurane and the muscles were dissected. Isolated muscles were frozen in a beaker filled with a slurry of isopentane at –80°C and stored in a deep freezer until further analysis. Frozen muscles were sectioned using a cryostat (Leica CM3050S) at 10 μm thickness. Cryosections were used for H&E staining and immunofluorescence analysis. For immunofluorescence, the sections were fixed with ice-cold methanol for 10 min, permeabilized with 0.25% Triton X-100 for 15 min, and blocked with 1% bovine serum albumin (BSA)/Tris-buffered saline with Tween (TBST) solution for 30 min at room temperature. The sections were incubated overnight at 4°C with primary antibodies (anti-V5, anti-laminin, anti-TOM20, streptavidin-Alexa Fluor 647) diluted in 1% BSA/TBST solution. After washing with TBST solution, the sections were incubated at room temperature for 1 h with the secondary antibody diluted in 1% BSA/TBST solution. Next, the stained sections were incubated with Hoechst 33342 solution for 5 min at room temperature and then mounted with anti-faded fluorescence mounting medium. All images were acquired on a Zeiss LSM 880 confocal microscope.

### APEX2 electron microscopy (MTS-V5-APEX2 and RTN4IP1-V5-APEX2)

To observe the DAB-stained mitochondria in the muscle tissues of MAX Tg mice, the dissected tissues were fixed with 2.5% glutaraldehyde and 2% paraformaldehyde in 0.1 M cacodylate solution (pH 7.0) for 1 h at 4°. After washing, 20 mM glycine solution was used to quench the unreacted aldehyde. DAB staining was performed for approximately 40 min until a light brown stain was visible under a stereomicroscope. DAB-stained tissues were post-fixed with 2% osmium tetroxide in distilled water for 60 min at 4°C and en bloc in 1% uranyl acetate overnight and dehydrated with a graded acetone series. The samples were then embedded with an Embed-812 embedding kit and polymerized in oven at 60 °C. The polymerized samples were sectioned (60 nm) with an ultramicrotome (UC7; Leica Microsystems, Germany), and the sections were mounted on copper slot grids with a specimen support film. Sections were stained with uranyless and lead citrate, and then observed on a Tecnai G2 transmission electron microscope (ThermoFisher, USA).

To visualize the subcellular localization of the transiently expressed RTN4IP1-V5-APEX2, HEK293T cells were cultured in 35 mm glass grid-bottomed culture dishes (MatTek Life Sciences, MA, USA) to 30– 40% confluency. The cells were then transfected with RTN4IP1-V5-APEX2 using Lipofectamine 2000. The next day, the cells were fixed with 2.5% glutaraldehyde and 2% paraformaldehyde in 0.1 M cacodylate solution (pH 7.0) for 1 h at 4°C. After washing, 20 mM glycine solution was used to quench unreacted aldehyde. DAB staining was performed for approximately 20−40 min until a light brown stain was visible under an inverted light microscope. DAB-stained cells were post-fixed with 2% osmium tetroxide in distilled water for 30 min at 4°C and dehydrated with a graded ethanol series. The samples were then embedded with the Embed-812 embedding kit and polymerized in an oven at 60°C. The polymerized samples were sectioned (60 nm) with an ultramicrotome (UC7; Leica Microsystems, Germany), and the sections were mounted on copper slot grids with a specimen support film. Sections were stained with uranyless and lead citrate, and were then viewed on a Tecnai G2 transmission electron microscope (ThermoFisher, USA).

### Electron microscopy of control and Rtn4IP1-KO C2C12 cells

Cells were grown in 35 mm glass-bottomed culture dishes to 50%–60% confluency. Cells were fixed with 2 ml of the fixative solution containing 2% paraformaldehyde and 2.5% of glutaraldehyde diluted in 0.1 M sodium cacodylate buffer. After washing, the cells were post-fixed in 2% osmium tetroxide (OsO_4_) containing 1.5% potassium ferrocyanide for 1 h at 4°C. The fixed cells were dehydrated using an ethanol series (50%, 60%, 70%, 80%, 90%, and 100%) for 10 min at each concentration and infiltrated with an embedding medium. After embedment, 60 nm sections were cut horizontally to the plane of the block (UC7; Leica Microsystems, Germany) and were mounted on copper slot grids with a specimen support film. The sections were then double-stained with uranyless and lead citrate, and observed using a Tecnai G2 transmission electron microscope at 120 kV in (ThermoFisher, USA).

### Immunofluorescence and confocal microscopy of HEK293T and C2C12 cells

To visualize the subcellular localization of the transiently expressed RTN4IP1-V5-APEX2, HEK293T cells were plated on coverslips (thickness no. 1.5, radius 18 mm). The cells were fixed in 4% paraformaldehyde and permeabilized with cold methanol for 5 min at –20°C, followed by washing with Dulbecco’s phosphate-buffered saline (DPBS) and blocking for 1 h with 2% BSA in DPBS at room temperature. Immunolabeling was conducted in blocking solution with appropriately diluted antibodies (anti-V5 and anti-TOM20) and Alexa Fluor-labeled secondary antibodies (anti-streptavidin–Alexa Fluor 647, mouse anti-Alexa Fluor 488, and rabbit anti-Alexa Fluor 568) with extensive washes. Immunofluorescence images were obtained and analyzed on an SP8 X Leica microscope (NICEM, Seoul National University, Seoul, Republic of Korea) with an objective lens (HC PL APO 100×/1.40 OIL), white light laser (470–670 nm, 1 nm tunable laser), and HyD detector, which was controlled with LAS X software.

### Recombinant RTN4IP1 expression and purification

The DNA fragments encoding WT RTN4IP1 were amplified by PCR and cloned into a modified pET-21a vector. For protein production, the plasmids were transformed into *Escherichia coli* BL21 (DE3) cells. Protein expression was induced with 0.3 mM isopropyl-β-D-1-thiogalactopyranoside (IPTG) when the cells reached an absorbance of 0.6 at an optical density of 600 nm; culturing was continued at 18°C for 18 h. The cells were harvested using centrifugation at 5000 ×*g* for 10 min, and resuspended in lysis buffer (25 mM sodium phosphate pH 7.8, 400 mM sodium chloride, and 10 mM imidazole). After cell lysis by sonication, lysed cells were clarified using centrifugation for 30 min at 10,000 ×*g*. The supernatant was applied onto an Ni2+-IMAC affinity column equilibrated with the binding buffer consisting of 25 mM sodium phosphate pH 7.8, 400 mM NaCl, and 10 mM imidazole. The proteins were eluted with binding buffer supplemented with 400 mM imidazole. The proteins were purified using Amicon filters (Milipore) and eluted in buffer with PBS. The protein solution was concentrated to approximately 5 mg/mL and flash-frozen in liquid nitrogen for storage.

### RTN4IP1 oxidoreductase activity

Oxidoreductase activity of RTN4IP1 was determined by measuring the absorption of 2,6-dichlorophenolindophenol (DCPIP; 50 μM) spectrophotometrically at 600 nm using the SpectraMax® i3x Multi-Mode Microplate Reader (Molecular Devices). Absorption was measured after adding purified RTN4IP1 (100 nM) protein or BSA (100 nM) to a reaction mixture containing 100 mM Tris·HCl, 0.01% Tween 20, and 200 μM NADPH (pH 7.4).

### Construction of the stably expressed RTN4IP1-V5-TurboID cell line

Flp-In™ T-Rex™ 293 cells were cultured in DMEM (Gibco) supplemented with 10% FBS, 2 mM L-glutamine, 50 units/mL penicillin, and 50 μg/mL streptomycin at 37°C under 5% CO2. Cells were grown in a T25 flask. Stable cell lines were first generated by transfection with the pcDNA5 expression construct plasmid expressing RTN4IP1-V5-TurboID or Matrix-V5-TurboID. Cells were transfected at 60–80% confluence using 6 μL of PEI transfection reagent and 2000 ng plasmid per 6-well cell culture plate. After 24 h, the cells were split into a 90 mm cell culture dish (SPL, 11090) with hygromycin (2 μg/mL). Media containing hygromycin were changed every 3–4 days. After 2–3 weeks, 3–4 colonies were selected and transferred to a 24-well plate.

The cells were continuously split into larger plates, and a cell stock was prepared. After splitting the cells into a 6-well plate, separate samples were prepared for expression detection. RTN4IP1-V5-TurboID or Matrix-V5-TurboID expression was induced by 5 ng/mL doxycycline.

### Construction of the Rtn4ip1-KO C2C12 cell line

The CRISPR/Cas9 technique was used to generate knockout cell lines. Single-guide RNAs (sgRNAs) were designed using the CRISPR RGEN Tools website (http://www.rgenome.net); the sequence 5′-GGAAGCGGTCGAAAGATAAA-3′ was used as a non-target control, while 5′-TCTGCCATAAACAAGGTTGG-3′ was used to target exon 4 of the mouse *Rtn4ip1* gene. Each sgRNA was cloned into lentiCRISPRv2. Production and transduction of CRISPR lentivirus was performed as previously described ^67^. Puromycin (2 μg/mL) was used to select knockout cells.

### Measurement of Coenzyme Q

Coenzymes Q9 and Q10, and their reduced forms, were determined by liquid chromatography-triple quadrupole mass spectrometry (LC-MS/MS) as previously described, with modifications ^68^. Briefly, freeze-dried cells were extracted with 300 μL methanol using a mixer mill (MM 440, Qiagen, Retsch Haan, Germany) at a frequency of 30 s with three repetitions. Extracts were placed on ice for 15 min and then centrifuged at 12,700 rpm at 4°C for 10 min. The supernatants were transferred to a clean tube. Remaining precipitate was re-extracted with methanol. The pooled supernatants were concentrated under a nitrogen stream, reconstituted with 100 μL methanol, and then subjected to LC-MS/MS. All analyses were performed using a Shimadzu LCMS 8060 MS/MS combined with a Nexera X2 LC system (Shimadzu, Kyoto, Japan) equipped with an electrospray ionization interface. Analytes were separated on a Luna C18 column (30 × 2 mm, 3 μm, 100 Å; Phenomenex, Torrance, CA, USA). The mobile phase consisted of 2 mM ammonium acetate in 100% methanol, and the flow rate was 300 μL/min under isocratic elution conditions. Electrospray ionization was operated in positive-ion mode at 4000 V. The optimum operating conditions were determined as follows: vaporizer temperature, 300°C; capillary temperature, 350°C; collision gas (argon) pressure, 1.5 mTorr.

Quantitation was conducted in selected reaction monitoring (SRM) modes with the precursor to product ion transition for each analyte (**Supplementary information**). The appropriate retention times for coenzyme Q10, Q10 reduced form, Q9, and Q9 reduced form were 8.5, 4.8, 5.3, and 3.2 min, respectively. The lower limits of quantification for coenzyme Q10, Q10 reduced form, Q9, and Q9 reduced form were 20 ng/mL, respectively. The interassay precision for all analytes was less than 9.4%.

### Immunofluorescence imaging of oxidized DNA

To visualize the 8-oxo guanosine, control or *Rtn4ip1*-KO C2C12 cells were plated on coverslips (thickness no. 1.5, radius 18 mm). The cells were fixed and permeabilized with cold methanol for 30 min at –20°C, followed by washing with DPBS. To eliminate RNA, the cells were treated with RNase solution for 1 h at 37°C. After washing three times with DPBS, nuclear DNA was denatured with 2 N HCl for 10 min. After washing with DPBS three times again, the cells were blocked for 1 h with 2% BSA in DPBS at room temperature. Immunolabeling was conducted in blocking solution with appropriately diluted anti-8-OHdG antibody and mouse anti-Alexa 568 with extensive washes. The cells were then counterstained with DAPI. Immunofluorescence images were obtained and analyzed on an SP8 X Leica microscope (NICEM) with an objective lens (HC PL APO 100×/1.40 OIL), white light laser (470–670 nm, 1 nm tunable laser), and HyD detector, which was controlled with LAS X software. Foci stained with anti-8-oxo-dG were counted using ImageJ software (National Institutes of Health).

### Measurement of membrane potential using TMRE

TMRE fluorescence analysis was conducted using flow cytometry and microplate reader. Control or *Rtn4ip1*-KO C2C12 cells were cultured in an incubator (37°C, 5% CO_2_). As a control experiment, the cells were treated with 200 μM of FCCP for 3 h.. For flow cytometry, the cells were harvested by trypsin treatment, and then resuspended in 0.5 mL DMEM supplemented with 5% FBS containing 200 nM of TMRE for 30 min at 37°C. Cellular fluorescence was measured using the Flow Activated Cell Sorter (FACS CantoII). Data were analyzed by BD FACSDiva™ Software. Microplate reader (Molecular Devices, Sunnyvale, CA, USA) measured fluorescence excitation at 520nm and emission at 580nm in a black 96-well culture plate with clear bottom.

### Oxygen Consumption Rate (OCR) measurement

Real-time measurements of OCR were performed using the Seahorse XFe96 Extracellular Flux Analyzer (Agilent) with Seahorse XF Cell Mito Stress Test Kit (103015-100). One day before the measurements, cells were plated at 1.2 × 10^4^ cells per well. One hour before the measurements, the cell culture medium was changed to pre-warmed DMEM supplemented with 10 mM glucose, 1 mM pyruvate, and 2 mM glutamine. To test mitochondrial stress, 1.5 μM oligomycin, 2 μM FCCP, 0.5 μM rotenone, and 0.5 μM antimycin A were utilized according to the suggested protocol of the manufacturer. OCR was measured according to the manufacturer’s Cell Mito Stress Test protocol. For knockdown experiments, 4.0 × 10^4^ cells/mL were reverse-transfected with siRNA and seeded at 0.8 × 10^4^ cells per well two days before the measurements.

### APEX2-mediated in situ biotinylation reaction in live cells

All cell lines expressing APEX2 were incubated with 250 μM biotin phenol for 30 min, followed by treatment with 1 mM H_2_O_2_ and quenching with 1 M sodium azide, Trolox, and sodium ascorbate. The cells were then lysed with RIPA buffer containing 1X protease cocktail for immunoblotting or fixed in 4% paraformaldehyde for immunofluorescence.

### In situ biotinylation reaction in muscle tissues of MAX-Tg mice

The dissected tissues were placed in test tubes and incubated with DBP (500 μM) in PBS for 1 h. Subsequently, diluted H_2_O_2_ (20 mM) was added to each sample for a final concentration of 2 mM H_2_O_2_, and the tubes were gently agitated for 2 min. The reaction was then quenched by adding DPBS containing 10 mM Trolox, 20 mM sodium azide, and 20 mM sodium ascorbate to the tubes. The labeled tissues were homogenized using a bead beater. Homogenized tissues were lysed with 1 mL RIPA buffer containing 1X protease inhibitor cocktail. Each sample was immunoblotted with anti-V5 and streptavidin-HRP to detect the expression of the processed bait protein (MTS-V5-APEX2) and DBP-modified proteins by Matrix-APEX2, respectively. Line scan analysis was performed using Image J. After subtraction of the background intensity value, protein signals from the top to the bottom of the protein marker were placed on the x-axis while the signal intensity from the top to the bottom of each lane was placed on the y-axis.

### Biotin labeling and sample preparation for LC-MS/MS analysis

#### Biotin labeling and tissue lysis

The dissected muscle tissues were placed in test tubes and incubated with DBP (500 μM) in PBS for 1 h. Diluted H_2_O_2_ (20 mM) was then added to each sample for a final concentration of 2 mM H_2_O_2_, and then the tubes were gently agitated for 2 min. The reaction was then quenched by adding DPBS containing 10 mM Trolox, 20 mM sodium azide, and 20 mM sodium ascorbate to the tubes. The labeled tissues were homogenized using a bead beater. Homogenized tissues were lysed with 1 mL lysis buffer (4% sodium dodecyl sulfate in 1X TBS containing 1X protease inhibitor cocktail). For clarifying, the lysates were ultrasonicated (Bioruptor) for 15 min; each step was performed on ice or at 4°C in a cold room.

#### Biotin labeling by RTN4IP1-V5-TurboID and cell lysis

For mass sampling, RTN4IP1-V5-TurboID and Matrix-V5-TurboID stable cells were grown in three T75 flasks to obtain triplicate samples. For transiently expressing constructs, cells at 70–80% confluence were treated with 5 ng/mL doxycycline. After 16 h, 50 μM biotin was added for 30 min and incubated at 37°C. After biotin labeling, the cells were washed three to four times with cold DPBS and lysed with 1 mL lysis buffer. For clarifying, the lysates were ultrasonicated for 15 min in a cold room.

#### Digestion and enrichment of biotinylated peptides

Cold acetone (4 mL) stored at –20°C was mixed with the lysates and stored at –20°C for at least 2 h. These samples were centrifuged at 13,000 ×*g* for 10 min at 4°C and the supernatant was gently discarded. The pellet was resuspended in 500 μL of 8 M urea in 50 mM ammonium bicarbonate. The protein concentration was determined using a BCA assay, after which the protein samples were denatured at 650 rpm for 1 h at 37°C using a Thermomixer (Eppendorf). Sample reduction and alkylation were individually performed by adding 10 mM dithiothreitol and 40 mM iodoacetoamide, respectively, and incubating at 650 rpm for 1 h at 37°C with the Thermomixer. The samples were diluted 8-fold using 50 mM ABC, after which CaCl_2_ was added for a final concentration of 1 mM. Samples were digested using trypsin (50:1 w/w) at 650 rpm for 6–18 h at 37°C with the Thermomixer. Insoluble material was removed by centrifuging for 3 min at 10,000 ×*g*. The SA beads (200 μL) were first washed with 2 M urea in 1X TBS three to four times and then added to the samples. The mixture was then rotated for 1 h at room temperature, followed by washing the beads two to three times with 2 M urea in 50 mM ABC; the flow-through fraction was not discarded. After discarding the supernatant, the beads were washed with pure water and transferred to new tubes. After adding 100 μL 80%, 0.2% TFA, and 0.1% formic acid, the biotinylated peptides were heated at 60°C and mixed at 650 rpm. The supernatants without SA beads were transferred to new tubes. This elution step was repeated at least four times, and then the total elution fractions were dried for 5 h using a Speed-vac (Eppendorf). The samples were stored at –20°C before use in LC-MS/MS analyses.

### LC-MS/MS analysis of enriched biotinylated peptide samples

Long analytical capillary columns (100 cm × 75 μm i.d.) and dual fritted trap columns (2 cm × 150 μm i.d) were packed in-house with 3 μm Jupiter C18 particles with pore size of 300 Å (Phenomenex, Torrance). The long analytical column was placed in a column heater (Analytical Sales and Services) regulated to a temperature of 45°C. NanoAcquity UPLC system (Waters, Milford) with back-flush mode for trapping of samples was operated at a flow rate of 300 nL/min over 2 h with linear gradient ranging from 95% solvent A (H_2_O with 0.1% formic acid) to 40% of solvent B (acetonitrile with 0.1% formic acid). The enriched samples were analyzed on an Orbitrap Fusion Lumos mass spectrometer (Thermo Scientific) equipped with an in-house customized nanoelectrospray ion source with the following instrumental parameters: spray voltage, 2.2 kV; capillary temperature, 275 °C; and RF lens level, 30.0. The precursor ion scan was operated with scan range of 300 to 1500 m/z; AGC target of 5e5; maximum injection time of 50 ms; and resolution of 120,000 at 200 m/z. The MS/MS scan was performed with isolation width of 1.4 Th; HCD with 30% collision energy; AGC target of 1e5; maximum injection time of 200 ms; and resolution of 30,000 at 200 m/z.

### LC-MS/MS data processing and identification of biotinylated peptides

All MS/MS data were searched by MaxQuant (version 1.6.2.3) with Andromeda search engine at 10 ppm precursor ion mass tolerance against the UniProt mouse reference proteome database (55,152 entries) or the SwissProt Homo sapiens proteome database (20,199 entries), according to sample’s origin. The label free quantification (LFQ) and Match Between Runs were used with the following search parameters: semi-trypic digestion, fixed carbaminomethylation on cysteine, dynamic oxidation of methionine, protein N-terminal acetylation, and dynamic DBP labeling (delta monoisotopic mass: +331.1896) of tyrosine for APEX2 samples or dynamic biotinylation of lysine (delta monoisotopic mass: +226.07759 Da) for TurboID samples. Less than 1% of false discovery rate was obtained at unique labeled peptide-level and unique labeled protein-level.

LFQ intensity values were log-transformed for further analysis and missing values were filled by imputed values representing a normal distribution around the detection limit. To impute the missing value, first, the intensity distribution of mean and standard deviation was determined, then for imputation values, new distribution based on Gaussian distribution with a downshift of 1.8 and width of 0.3 standard deviations was created for total matrix.

### Determination of mitochondrial matrix targeting proteins with Mitofates probability scores

To obtain the subcellular localization information of our identified proteins, we refer to “subcellular localization” information in the Uniprot Database (www.uniprot.org, lastly modified in January, 2021). For the determination of mitochondrial matrix targeting proteins, Mitofates (http://mitf.cbrc.jp/MitoFates/cgi-bin/top.cgi) was used. Mitofates is a web-based prediction tool for N-terminal mitochondrial matrix targeting sequence (MTS) with probability score ^29^. Using this tool, proteins with MTS probability score above 0.1 were regarded as mitochondrial matrix proteins according to the minimum probability score of mitochondrial proteins with bona fide MTS such as UQCRB (Mitofates score=0.126) ^69^ and MCU (0.106) ^26^.

## Resource availability

### Lead Contact

Further information and requests for resources and reagents should be directed to and will be fulfilled by the lead contact, Hyun-Woo Rhee.

### Materials availability

Upon reasonable request, unique reagents utilized in this paper can be provided.

### Data availability

Upon reasonable request, unique reagents utilized in this paper can be provided. The mass spectrometry proteomics data have been deposited to the ProteomeXchange Consortium (http://proteomecentral.proteomexchange.org) via the PRIDE partner repository ^70^ with the dataset identifier PXD026793 (username; password: arAcld4N).

## Acknowledgements

This work was supported by the National Research Foundation of Korea (NRF-2019R1A2C3008463 to H.W.R, NRF-2018R1A2A3075389, NRF-2017K1A1A2013124 to J.M.S., and NRF-2019M3E5D3073104 to J.S.K.), Organelle Network Research Center (NRF-2017R1A5A1015366), and a grant from the Korea Health Industry Development Institute (KHIDI) funded by the Ministry of Health & Welfare and Ministry of Science and Information & Communication Technology (ICT), Republic of Korea (grant number HU20C0326). J.S.K. thanks to the support of the Institute for Basic Science (IBS-R008-D1) funded by the Ministry of Science and ICT of Korea and the New Faculty Startup Fund from Seoul National University. J.Y.M. and M.J. are supported by the Korea Brain Research Institute (KBRI) Basic Research Program funded by the Ministry of Science and ICT (21-BR-01-11). H.W.R. was supported by the Creative-Pioneering Researchers Program through Seoul National University. Electron microscopy data were acquired at the Brain Research Core Facilities of the KBRI.

## Author contributions

Conceptualization, I.P., K.E.K., J.M.S., H.W.R.; Investigation, all authors.; Supervision, J.S.K., J.M.S., and H.W.R; Writing, I.P., K.E.K., J.S.K., J.M.S., H.W.R.

## Extended data

**Extended Data Fig. 1. Experimental scheme to profile the mitochondrial matrix proteomes of mouse muscle tissues**.

**(a)** Scheme of LC-MS/MS analysis using MAX-Tg mice. **(b)** Volcano plot for the DBP-labeled proteome of each muscle tissues from WT mice (left) vs. MAX-Tg mice (right). Both samples were treated with the same amount of DBP and H_2_O_2_ prior to mass sampling. The cut-off for the mitochondrial matrix proteins was P < 0.05 and fold change (FC) > 2. See **Supplementary Table 3** for detailed information.

**Extended Data Fig. 2 (Related to Fig. 3b**). **Asymmetrical DBP-labeled mitochondrial matrix proteins detected by MTS-V5-APEX2 in the TA and heart muscles of MAX-Tg mice**.

Heart and TA muscle-enriched proteins are in light blue and pink, respectively. All proteins are color-coded to reflect the fold change in the average intensity between the TA muscle and heart. Annotations with function and complex are based on information from UNIPROT and CORUM. See **Supplementary Table 5** for detailed information.

**Extended Data Fig. 3 (Related to Fig. 5b). Functional clustering with the RTN4IP1 interactome using STRING analysis**.

Molecular interaction network of 32 significant proteins biotinylated by RTN4IP1-TurboID (RTN4IP1 interactome) were analyzed by STRING analysis ^71^ (https://string-db.org/). The clustered molecular functions are marked with different colors: red, coenzyme Q biosynthesis; blue, mitochondrial complex I assembly; green, mitochondrial translation elongation and termination; yellow, protein targeting to the mitochondrion.

**Extended Data Fig. 4 (Related to Fig. 6a, b). Additional TEM images of control (upper panel) and *Rtn4ip1-*KO (lower panel) C2C12 cells**.

**Extended Data Fig. 5 Structural similarity of RTN4IP1 orthologs**.

**(a)** Structural similarity between crystal structure of RTN4IP1 (Uniprot ID: Q8WWV3, PDB ID: 2VN8) and the predicted structure in AlphfaFold database (AF-Q8WWV3-F1, https://alphafold.ebi.ac.uk/) **(b)** Structural similarity between human RTN4IP1 and RTN4IP1 orthologs: CG17221 (Drosophila, AF-Q8IPZ3-F1), **(c)** Rad-8 (C. elegans, AF-P28625-F1), **(d)** Yim1p (Yeast, AF-P28625-F1), and **(e)** qorA(E. coli, AF-P28304-F1) from AlphaFold database

**Supplementary Table 1. DBP-labeled peptides and proteins in MAX-Tg**

**Supplementary Table 2. DBP-labeled proteins by MTS-APEX2 in HEK293T and in muscle tisseus_related with Fig. 2a**

**Supplementary Table 3. DBP-labeled proteins in wild type and in MAX-Tg_related with Extended Data Fig. 1b**

**Supplementary Table 4. T-Test for DBP-labeled proteins by MTS-APEX2 in TA and in HEK293T_related with Fig. 2e, f**

**Supplementary Table 5. Tissue-specific DBP-labeled proteins in MAX-Tg_related with Fig. 3b-d and Fig. 2**

**Supplementary Table 6. RTN4IP1 interactome_related with Fig. 5a-c**

**Supplementary Table 7. Construct Information**

**Supplementary Table 8. SRM transitions used for quantification of coenzyme Q10 (oxidized form), coenzyme Q10 (reduced form), coenzyme Q9 (oxidized form), and coenzyme Q9 (reduced form)**

## REFERENCES

1. Uhlen, M. et al. Tissue-based map of the human proteome. Science 347, 1260419 (2015).

2. Uhlen, M. et al. A genome-wide transcriptomic analysis of protein-coding genes in human blood cells. Science 366, eaax9198 (2019).

3. Zick, M., Rabl, R. & Reichert, A.S. Cristae formation-linking ultrastructure and function of mitochondria. Biochimica et biophysica acta 1793, 5–19 (2009).

4. Pette, D. & Vrbová, G. What does chronic electrical stimulation teach us about muscle plasticity. Muscle and Nerve 22, 666–677 (1999).

5. Johannsen, D.L. & Ravussin, E. Can Increased Muscle ROS Scavenging Keep Older Animals Young and Metabolically Fit? Cell Metabolism 12, 557–558 (2010).

6. Williams, E.G. et al. Quantifying and Localizing the Mitochondrial Proteome Across Five Tissues in A Mouse Population. Mol Cell Proteomics 17, 1766–1777 (2018).

7. Mootha, V.K. et al. Integrated analysis of protein composition, tissue diversity, and gene regulation in mouse mitochondria. Cell 115, 629–640 (2003).

8. Bayraktar, E.C. et al. MITO-Tag Mice enable rapid isolation and multimodal profiling of mitochondria from specific cell types in vivo. Proc Natl Acad Sci U S A 116, 303–312 (2019).

9. Busch, J.D. et al. MitoRibo-Tag Mice Provide a Tool for In Vivo Studies of Mitoribosome Composition. Cell reports 29, 1728–1738 e1729 (2019).

10. Yoo, C.-M. & Rhee, H.-W. APEX, a Master Key To Resolve Membrane Topology in Live Cells. Biochemistry 59, 250–259 (2020).

11. Lee, S.-Y. et al. Architecture Mapping of the Inner Mitochondrial Membrane Proteome by Chemical Tools in Live Cells. J Am Chem Soc 139, 3651–3662 (2017).

12. Silva, J. et al. EXD2 governs germ stem cell homeostasis and lifespan by promoting mitoribosome integrity and translation. Nature Cell Biology 20, 162–174 (2018).

13. Park, J. et al. The structure of human EXD2 reveals a chimeric 3′ to 5′ exonuclease domain that discriminates substrates via metal coordination. Nucleic Acids Research 47, 7078–7093 (2019).

14. Rhee, H.-W. et al. Proteomic mapping of mitochondria in living cells via spatially restricted enzymatic tagging. Science (New York, N.Y.) 339, 1328–1331 (2013).

15. Lobingier, B.T. et al. An Approach to Spatiotemporally Resolve Protein Interaction Networks in Living Cells. Cell 169, 350-360.e312 (2017).

16. Lam, S.S. et al. Directed evolution of APEX2 for electron microscopy and proximity labeling. Nature Methods 12, 51–54 (2015).

17. Hung, V. et al. Proteomic mapping of the human mitochondrial intermembrane space in live cells via ratiometric APEX tagging. Molecular cell 55, 332–341 (2014).

18. Udeshi, N.D. et al. Antibodies to biotin enable large-scale detection of biotinylation sites on proteins. Nature Methods 14, 1167–1170 (2017).

19. Kwak, C. et al. Contact-ID, a tool for profiling organelle contact sites, reveals regulatory proteins of mitochondrial-associated membrane formation. Proceedings of the National Academy of Sciences 117, 12109 (2020).

20. Cho, K.F. et al. Split-TurboID enables contact-dependent proximity labeling in cells. Proceedings of the National Academy of Sciences 117, 12143 (2020).

21. Calvo, S.E., Clauser, K.R. & Mootha, V.K. MitoCarta2.0: an updated inventory of mammalian mitochondrial proteins. Nucleic Acids Research 44, D1251–D1257 (2016).

22. Rath, S. et al. MitoCarta3.0: an updated mitochondrial proteome now with sub-organelle localization and pathway annotations. Nucleic Acids Research 49, D1541–D1547 (2021).

23. Pagliarini, D.J. et al. A mitochondrial protein compendium elucidates complex I disease biology. Cell 134, 112–123 (2008).

24. Rahbani, J.F. et al. Creatine kinase B controls futile creatine cycling in thermogenic fat. Nature 590, 480–485 (2021).

25. Sun, Y. et al. Mitochondrial TNAP controls thermogenesis by hydrolysis of phosphocreatine. Nature 593, 580–585 (2021).

26. Martell, J.D. et al. Engineered ascorbate peroxidase as a genetically encoded reporter for electron microscopy. Nature biotechnology 30, 1143–1148 (2012).

27. Picard, M., White, K. & Turnbull, D.M. Mitochondrial morphology, topology, and membrane interactions in skeletal muscle: a quantitative three-dimensional electron microscopy study. Journal of Applied Physiology 114, 161–171 (2012).

28. Palmer, J.W., Tandler, B. & Hoppel, C.L. Biochemical properties of subsarcolemmal and interfibrillar mitochondria isolated from rat cardiac muscle. The Journal of biological chemistry 252, 8731–8739 (1977).

29. Fukasawa, Y. et al. MitoFates: Improved Prediction of Mitochondrial Targeting Sequences and Their Cleavage Sites*. Molecular & Cellular Proteomics 14, 1113–1126 (2015).

30. Ghezzi, D. & Zeviani, M. Assembly Factors of Human Mitochondrial Respiratory Chain Complexes: Physiology and Pathophysiology, in Mitochondrial Oxidative Phosphorylation: Nuclear-Encoded Genes, Enzyme Regulation, and Pathophysiology. (ed.B. Kadenbach) 65–106 (Springer New York, New York, NY; 2012).

31. Borna, N.N. et al. Mitochondrial ribosomal protein PTCD3 mutations cause oxidative phosphorylation defects with Leigh syndrome. neurogenetics 20, 9–25 (2019).

32. Newman, A.C. & Maddocks, O.D.K. One-carbon metabolism in cancer. British Journal of Cancer 116, 1499–1504 (2017).

33. Altman, B.J., Stine, Z.E. & Dang, C.V. From Krebs to clinic: glutamine metabolism to cancer therapy. Nature reviews. Cancer 16, 619–634 (2016).

34. Staron, R.S. et al. Fiber type composition of four hindlimb muscles of adult Fisher 344 rats. Histochemistry and Cell Biology 111, 117–123 (1999).

35. Lopaschuk, G.D., Ussher, J.R., Folmes, C.D.L., Jaswal, J.S. & Stanley, W.C. Myocardial Fatty Acid Metabolism in Health and Disease. Physiological Reviews 90, 207–258 (2010).

36. Oizel, K. et al. D-2-Hydroxyglutarate does not mimic all the IDH mutation effects, in particular the reduced etoposide-triggered apoptosis mediated by an alteration in mitochondrial NADH. Cell Death & Disease 6, e1704–e1704 (2015).

37. Angebault, C. et al. Recessive Mutations in RTN4IP1 Cause Isolated and Syndromic Optic Neuropathies. Am J Hum Genet 97, 754–760 (2015).

38. Charif, M. et al. Neurologic Phenotypes Associated With Mutations in RTN4IP1 (OPA10) in Children and Young Adults. JAMA Neurol 75, 105–113 (2018).

39. Thorn, J.M., Barton, J.D., Dixon, N.E., Ollis, D.L. & Edwards, K.J. Crystal structure of Escherichia coli QOR quinone oxidoreductase complexed with NADPH. Journal of molecular biology 249, 785–799 (1995).

40. Taneja, B. & Mande, S.C. Conserved structural features and sequence patterns in the GroES fold family. Protein engineering 12, 815–818 (1999).

41. Murzin, A.G. Structural classification of proteins: new superfamilies. Current opinion in structural biology 6, 386–394 (1996).

42. Sillitoe, I. et al. CATH: expanding the horizons of structure-based functional annotations for genome sequences. Nucleic Acids Research 47, D280–D284 (2019).

43. Bongard, R.D., Lindemer, B.J., Krenz, G.S. & Merker, M.P. Preferential utilization of NADPH as the endogenous electron donor for NAD(P)H:quinone oxidoreductase 1 (NQO1) in intact pulmonary arterial endothelial cells. Free Radical Biology and Medicine 46, 25–32 (2009).

44. Bongard, R.D. et al. Characterization of the threshold for NAD(P)H:quinone oxidoreductase activity in intact sulforaphane-treated pulmonary arterial endothelial cells. Free Radic Biol Med 50, 953–962 (2011).

45. Merker, M.P., Audi, S.H., Bongard, R.D., Lindemer, B.J. & Krenz, G.S. Influence of pulmonary arterial endothelial cells on quinone redox status: effect of hyperoxia-induced NAD(P)H:quinone oxidoreductase 1. American Journal of Physiology-Lung Cellular and Molecular Physiology 290, L607–L619 (2006).

46. Branon, T.C. et al. Efficient proximity labeling in living cells and organisms with TurboID. Nature Biotechnology 36, 880–887 (2018).

47. Lee, S.Y., Seo, J.K. & Rhee, H.W. Direct Identification of Biotinylated Proteins from Proximity Labeling (Spot-BioID). Methods Mol Biol 2008, 97–105 (2019).

48. Stefely, J.A. & Pagliarini, D.J. Biochemistry of Mitochondrial Coenzyme Q Biosynthesis. Trends in biochemical sciences 42, 824–843 (2017).

49. Stefely, J.A. et al. Cerebellar Ataxia and Coenzyme Q Deficiency through Loss of Unorthodox Kinase Activity. Molecular cell 63, 608–620 (2016).

50. Bersuker, K. et al. The CoQ oxidoreductase FSP1 acts parallel to GPX4 to inhibit ferroptosis. Nature 575, 688–692 (2019).

51. Mellors, A. & Tappel, A.L. The inhibition of mitochondrial peroxidation by ubiquinone and ubiquinol. The Journal of biological chemistry 241, 4353–4356 (1966).

52. Doll, S. et al. ACSL4 dictates ferroptosis sensitivity by shaping cellular lipid composition. Nat. Chem. Biol. 13, 91–98 (2017).

53. Friedmann Angeli, J.P. et al. Inactivation of the ferroptosis regulator Gpx4 triggers acute renal failure in mice. Nature Cell Biology 16, 1180–1191 (2014).

54. Lewerenz, J., Ates, G., Methner, A., Conrad, M. & Maher, P. Oxytosis/Ferroptosis-(Re-) Emerging Roles for Oxidative Stress-Dependent Non-apoptotic Cell Death in Diseases of the Central Nervous System. Front Neurosci 12, 214 (2018).

55. Höglinger, D. et al. NPC1 regulates ER contacts with endocytic organelles to mediate cholesterol egress. Nature Communications 10, 4276 (2019).

56. Daigle, T.L. et al. A Suite of Transgenic Driver and Reporter Mouse Lines with Enhanced Brain-Cell-Type Targeting and Functionality. Cell 174, 465-480.e422 (2018).

57. Hu, W.-H. et al. Identification and characterization of a novel Nogo-interacting mitochondrial protein (NIMP). Journal of Neurochemistry 81, 36–45 (2002).

58. Dixon, Scott J. et al. Ferroptosis: An Iron-Dependent Form of Nonapoptotic Cell Death. Cell 149, 1060–1072 (2012).

59. Li, J. et al. Ferroptosis: past, present and future. Cell Death & Disease 11, 88 (2020).

60. Doll, S. et al. FSP1 is a glutathione-independent ferroptosis suppressor. Nature 575, 693–698 (2019).

61. Zou, X.H. et al. Whole Exome Sequencing Identifies Two Novel Mutations in the Reticulon 4-Interacting Protein 1 Gene in a Chinese Family with Autosomal Recessive Optic Neuropathies. Journal of molecular neuroscience : MN 68, 640–646 (2019).

62. Giacomini, T. et al. Optic Atrophy and Generalized Chorea in a Patient Harboring an OPA10/RTN4IP1 Pathogenic Variant. Neuropediatrics 51, 425–429 (2020).

63. D’Gama, A.M. et al. Exome sequencing identifies novel missense and deletion variants in RTN4IP1 associated with optic atrophy, global developmental delay, epilepsy, ataxia, and choreoathetosis. American Journal of Medical Genetics Part A 185, 203–207 (2021).

64. Fujii, M., Yasuda, K., Hartman, P.S., Ayusawa, D. & Ishii, N. A mutation in a mitochondrial dehydrogenase/reductase gene causes an increased sensitivity to oxidative stress and mitochondrial defects in the nematode Caenorhabditis elegans. Genes to cells : devoted to molecular & cellular mechanisms 16, 1022–1034 (2011).

65. Birrell, G.W. et al. Transcriptional response of Saccharomyces cerevisiae to DNA-damaging agents does not identify the genes that protect against these agents. Proc Natl Acad Sci U S A 99, 8778–8783 (2002).

66. Jumper, J. et al. Highly accurate protein structure prediction with AlphaFold. Nature 596, 583–589 (2021).

67. Joung, J. et al. Genome-scale CRISPR-Cas9 knockout and transcriptional activation screening. Nature protocols 12, 828–863 (2017).

68. Tang, Z. et al. Rapid assessment of the coenzyme Q10 redox state using ultrahigh performance liquid chromatography tandem mass spectrometry. The Analyst 139, 5600–5604 (2014).

69. Gu, J. et al. The architecture of the mammalian respirasome. Nature 537, 639–643 (2016).

70. Perez-Riverol, Y. et al. The PRIDE database and related tools and resources in 2019: improving support for quantification data. Nucleic Acids Res 47, D442–d450 (2019).

71. Szklarczyk, D. et al. STRING v11: protein-protein association networks with increased coverage, supporting functional discovery in genome-wide experimental datasets. Nucleic Acids Res 47, D607–d613 (2019).

