## Supplemental data figure and table for "In vivo mitochondrial matrix proteome profiling reveals RTN4IP1/OPA10 as an antioxidant NADPH oxidoreductase"

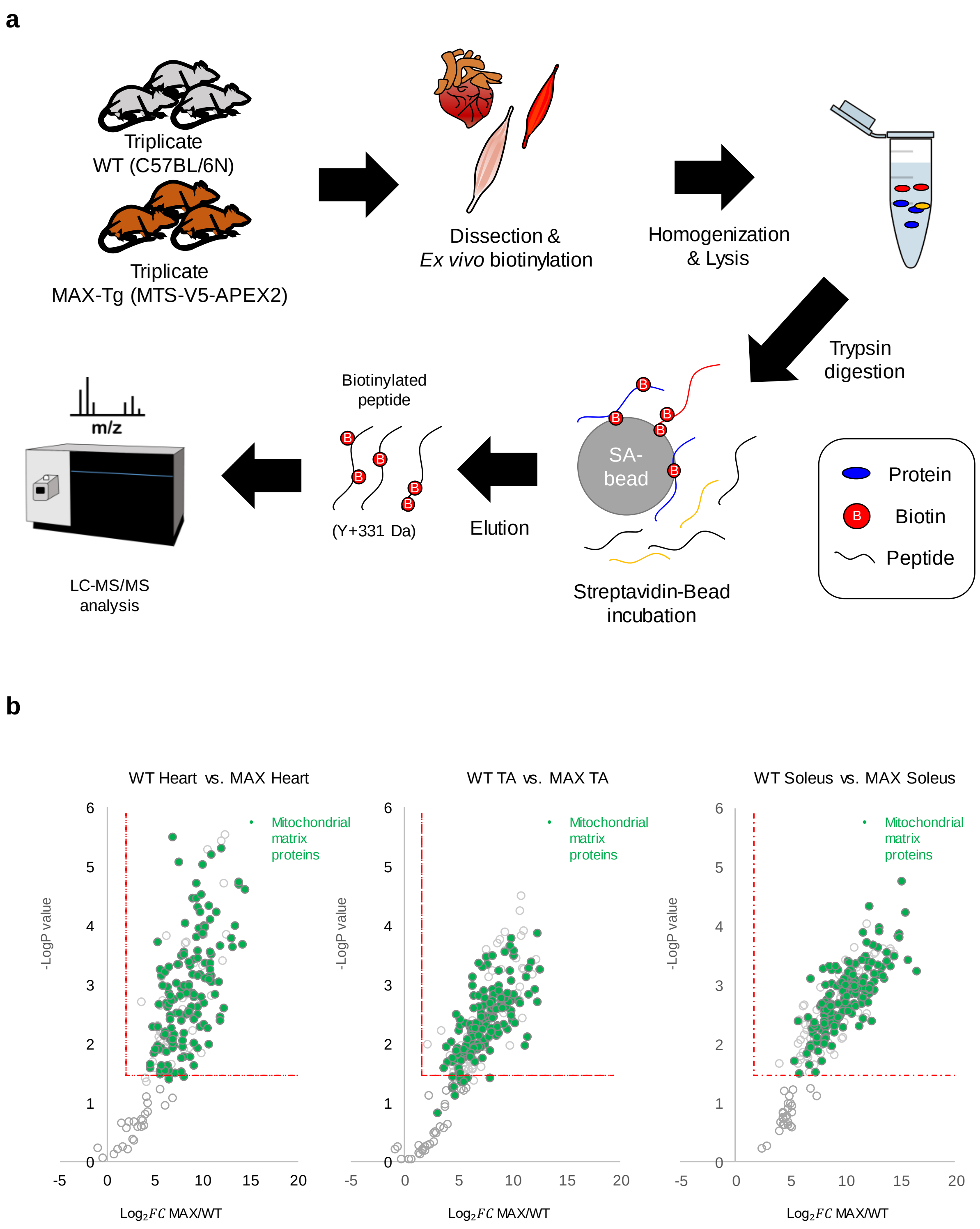
a
Triplicate
WT (C57BL/6N)
Triplicate
MAX-Tg (MTS-V5-APEX2)
Dissection &
*Ex vivo* biotinylation
Homogenization
& Lysis
Trypsin
digestion
SA-
bead
B
Streptavidin-Bead
incubation
Protein
Biotin
Peptide
Elution
Biotinylated
peptide
(Y+331 Da)
m/z
LC-MS/MS
analysis
b
WT Heart vs. MAX Heart
WT TA vs. MAX TA
WT Soleus vs. MAX Soleus
Mitochondrial
matrix
proteins
-LogP value
Log$_2$ $FC$ MAX/WT

**Extended Data Fig. 1: Experimental scheme to profile the mitochondrial matrix proteomes of mouse muscle tissues.**

**(a)** Scheme of LC-MS/MS analysis using MAX-Tg mice. **(b)** Volcano plot for the DBP-labeled proteome of each muscle tissues from WT mice (left) vs. MAX-Tg mice (right). Both samples were treated with the same amount of DBP and $H_2O_2$ prior to mass sampling. The cut-off for the mitochondrial matrix proteins was $P < 0.05$ and fold change (FC) > 2. See Table S3 for detailed information.

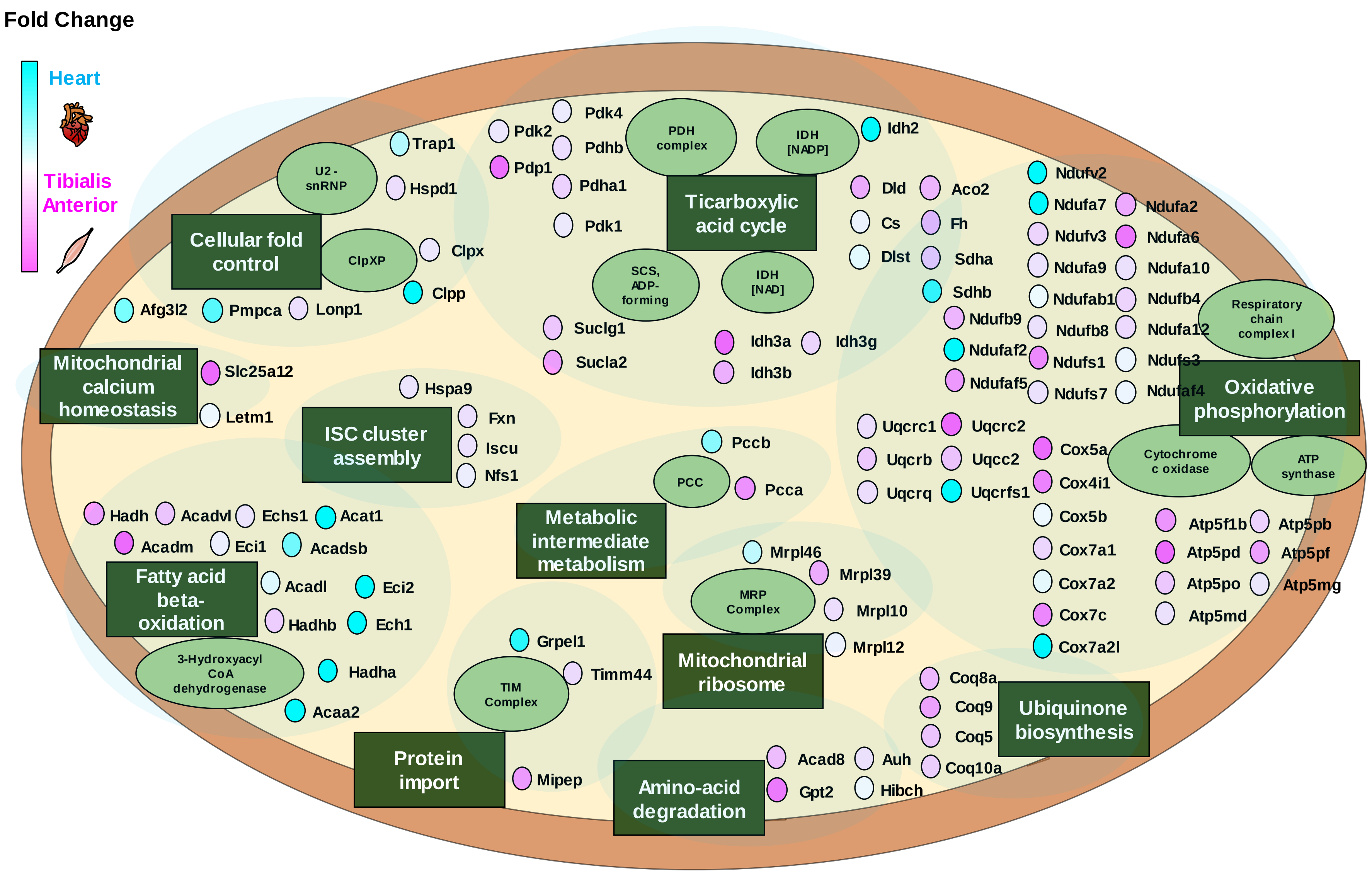

Fold Change
Heart
Tibialis Anterior
Cellular fold control
U2-snRNP
ClpXP
Trap1
Hspd1
Clpx
Clpp
Afg3l2
Pmpca
Lonp1
Mitochondrial calcium homeostasis
Slc25a12
Letm1
ISC cluster assembly
Hspa9
Fxn
Iscu
Nfs1
Fatty acid beta-oxidation
Hadh
Acadvl
Echs1
Acat1
Acadm
Eci1
Acadsb
Acadl
Eci2
Hadhb
Ech1
3-Hydroxyacyl CoA dehydrogenase
Hadha
Acaa2
Protein import
TIM Complex
Grpel1
Timm44
Mipep
Metabolic intermediate metabolism
PCC
Pccb
Pcca
Mitochondrial ribosome
MRP Complex
Mrpl46
Mrpl39
Mrpl10
Mrpl12
Amino-acid degradation
Acad8
Auh
Gpt2
Hibch
Ticarboxylic acid cycle
PDH complex
IDH [NADP]
SCS, ADP-forming
IDH [NAD]
Pdk2
Pdk4
Pdhb
Pdp1
Pdha1
Pdk1
Suclg1
Sucla2
Idh3a
Idh3g
Idh3b
Idh2
Dld
Aco2
Cs
Fh
Dlst
Sdha
Sdhb
Oxidative phosphorylation
Respiratory chain complex I
Cytochrome c oxidase
ATP synthase
Ndufv2
Ndufa7
Ndufa2
Ndufv3
Ndufa6
Ndufa9
Ndufa10
Ndufab1
Ndufb4
Ndufb9
Ndufb8
Ndufa12
Ndufaf2
Ndufs1
Ndufs3
Ndufaf5
Ndufs7
Ndufaf4
Uqcrc1
Uqcrc2
Uqcrb
Uqcc2
Uqcrq
Uqcrfs1
Cox5a
Cox4i1
Cox5b
Cox7a1
Cox7a2
Cox7c
Cox7a2l
Atp5f1b
Atp5pb
Atp5pd
Atp5pf
Atp5po
Atp5mg
Atp5md
Ubiquinone biosynthesis
Coq8a
Coq9
Coq5
Coq10a

**Extended Data Fig. 2 (Related to Fig. 3b): Asymmetrical DBP-labeled mitochondrial matrix proteins detected by MTS-V5-APEX2 in the TA and heart muscles of MAX-Tg**

Heart and TA muscle-enriched proteins are in light blue and pink, respectively. All proteins are color-coded to reflect the fold change in the average intensity between the TA muscle and heart. Annotations with function and complex are based on information from UNIPROT and CORUM. See **Supplementary Table 5** for detailed information.

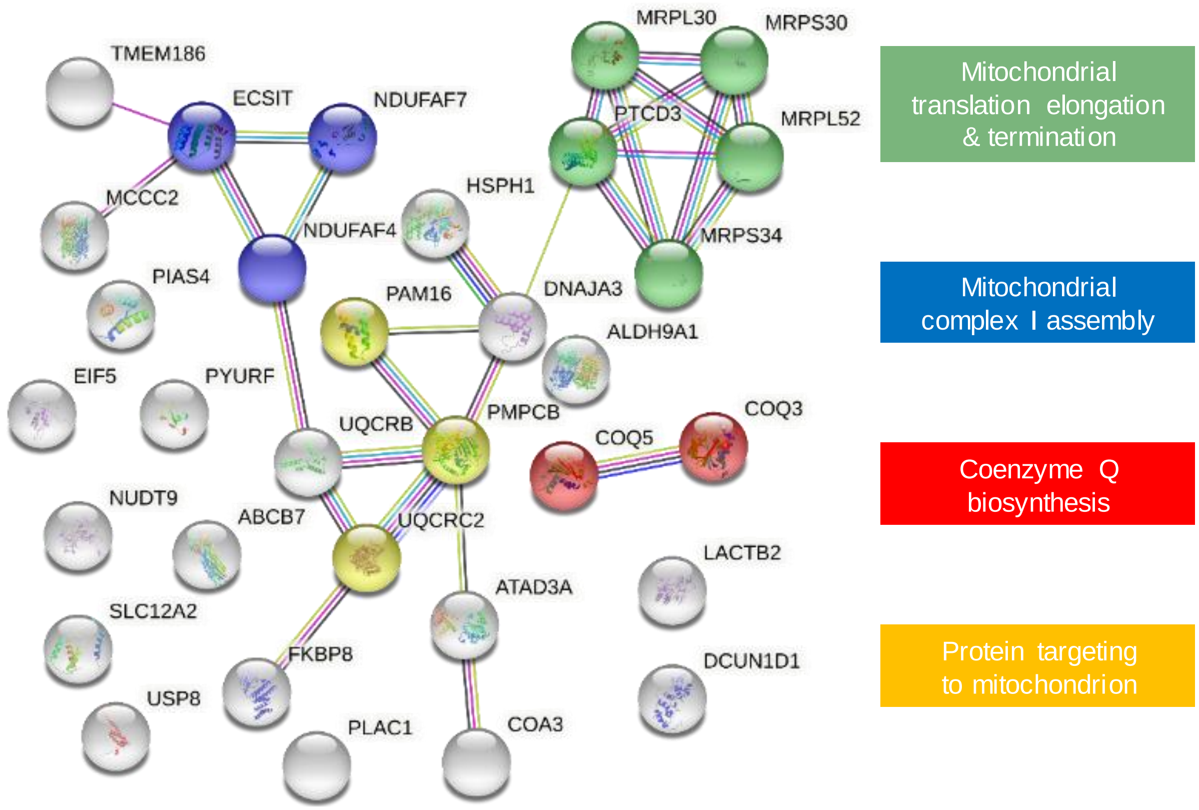


**Extended Fig. 3 (Related to Fig. 5b): Functional clustering with the RTN4IP1 interactome using STRING analysis**.

Molecular interaction network of 32 significant proteins biotinylated by RTN4IP1-TurboID (RTN4IP1 interactome) were analyzed by STRING analysis (https://string-db.org/). The clustered molecular functions are marked with different colors: red, coenzyme Q biosynthesis; blue, mitochondrial complex I assembly; green, mitochondrial translation elongation and termination; yellow, protein targeting to the mitochondrion.

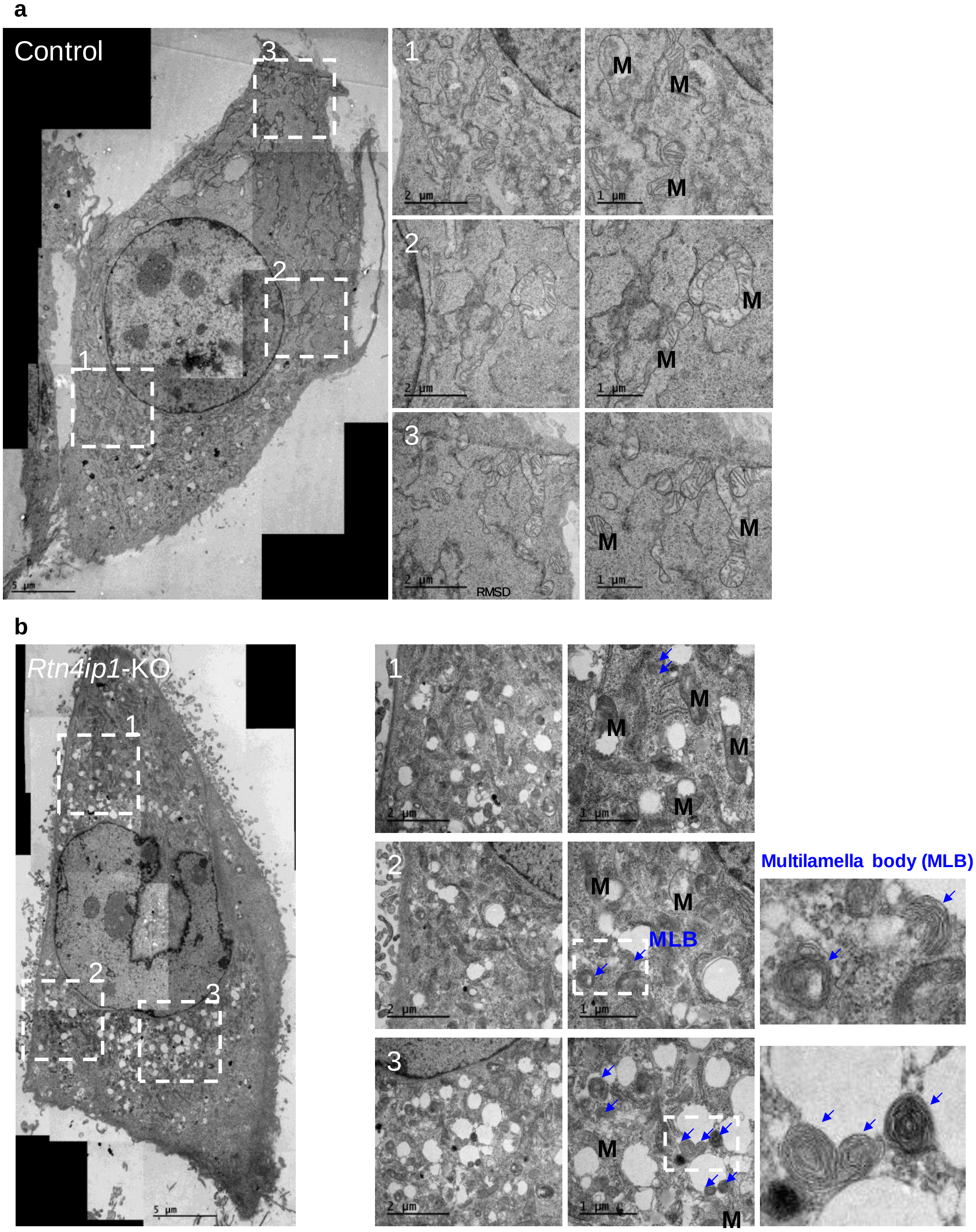


**Extended Data Fig. 4 (Related to Fig. 6a, b): Additional TEM images of control (upper panel) and *Rtn4ip1*-KO (lower panel) C2C12 cells.**

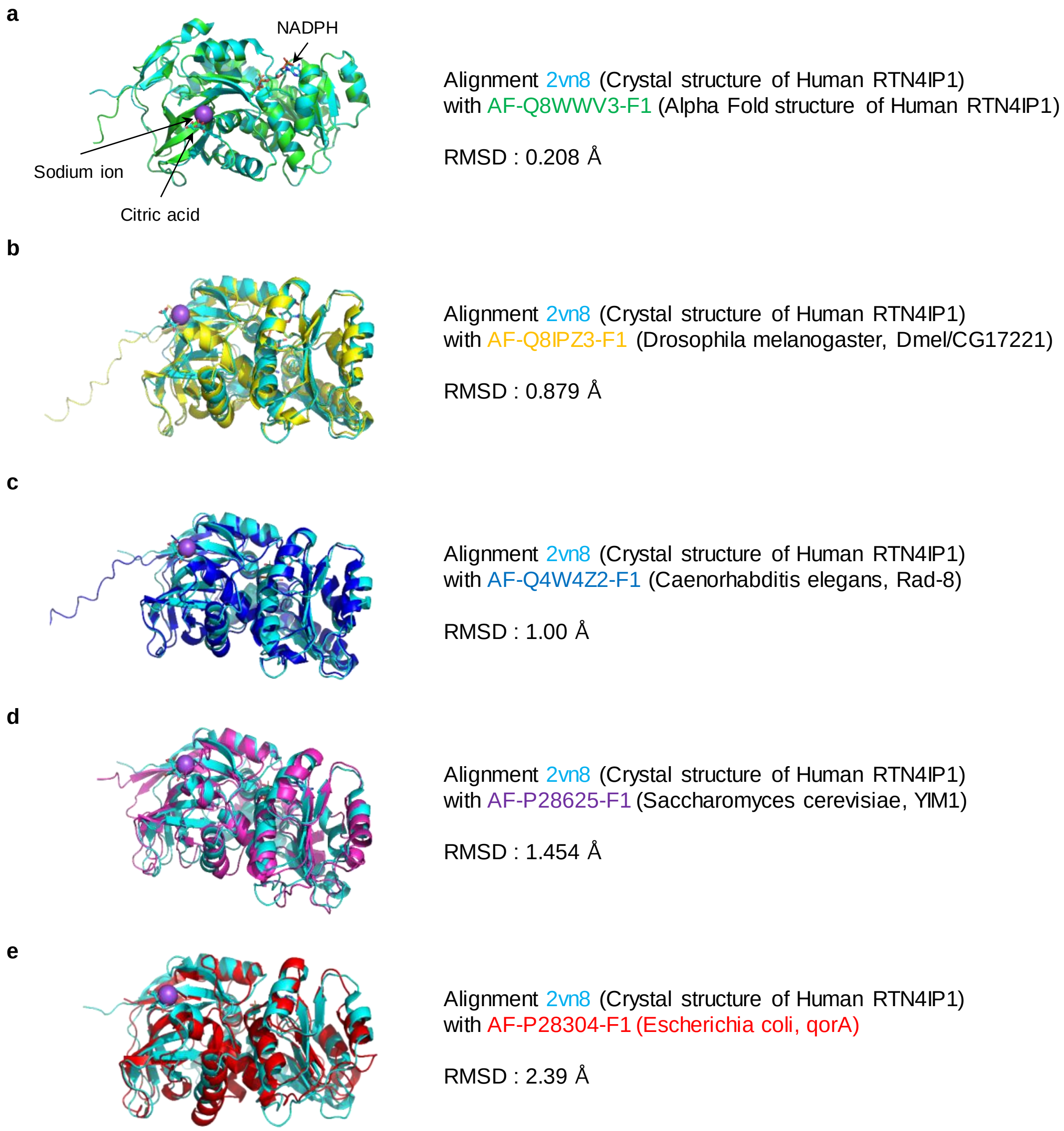


**Extended Data Fig. 5 Structural similarity of RTN4IP1 orthologs.**

**(a)** Structural similarity between crystal structure of RTN4IP1 (Uniprot ID: Q8WWV3, PDB ID: 2VN8) and the predicted structure in AlphfaFold database (AF-Q8WWV3-F1, https://alphafold.ebi.ac.uk/) **(b)** structural similarity between human RTN4IP1 and RTN4IP1 orthologs: CG17221 (Drosophila, AF-Q8IPZ3-F1), **(c)** Rad-8 (C. elegans, AF-P28625-F1), **(d)** Yim1p (Yeast, AF-P28625-F1), and **(e)** qorA(E. coli, AF-P28304-F1) from Alphfafold database

**Supplementary Table 1: DBP-labeled peptides and proteins in MAX-Tg**

**Supplementary Table 2: DBP-labeled proteins by MTS-APEX2 in HEK293T and in muscle tisseus_related with Fig. 2a**

**Supplementary Table 3: DBP-labeled proteins in wild type and in MAX-Tg_related with Extended Fig. 1b**

**Supplementary Table 4: T-Test for DBP-labeled proteins by MTS-APEX2 in TA and in HEK293T_related with Fig. 2e, f**

**Supplementary Table 5: Tissue-specific DBP-labeled proteins in MAX-Tg_related with Fig. 3b-d and Extended Data Fig. 2**

**Supplementary Table 6: RTN4IP1 interactome_related with Fig 5a-c**

**Supplementary Table 7: Construct Information**

| Name | Features | Promotor/Vector | Details |
|---|---|---|---|
| MTS-V5-APEX2_pCDNA5 | KpnI-MTS-BamHINheI-V5-APEX2-Stop-NotI | CMV/pCDNA5 | MTS-V5-APEX2 was a gift from Prof. Alice Ting (Addgene plasmid #72480) |
| RTN4IP1-V5-APEX2_pCDNA5 | AflII-RTN4IP1 NheI-V5-APEX2-Stop-NotI | CMV/pCDNA5 | NA |
| MTS(from RTN4IP1)-V5-APEX2_pCDNA5 | AflII-MTS(from RTN4IP1)-NheI-V5-APEX2-Stop-NotI | CMV/pCDNA5 | NA |
| RTN4IP1(Δ1-32aa)-V5-APEX2_pCDNA5 | AflII-RTN4IP1(Δ1-32aa)-NheI-V5-APEX2-Stop-NotI | CMV/pCDNA5 | NA |
| His6_MBP_TEV_ RTN4IP1(Δ1-32aa)_pET21a | NdeI-His6-NdeI-MBP-EcoRINheI-TEV cleavage site-RTN4IP1-Stop-XhoI | T7 /pET21a | NA |
| RTN4IP1-TurboID_pCDNA5 | AflII-RTN4IP1 KpnI-V5-TurboID-Stop-XhoI | CMV/pCDNA5 | NA |
| MTS-V5-TurboID_pCDNA5 | AflII-MTS-KpnI-V5-TurboID-Stop-XhoI | CMV/pCDNA5 | NA |

**Supplementary Table 8: SRM transitions used for quantification of coenzyme Q10 (oxidized form), coenzyme Q10 (reduced form), coenzyme Q9 (oxidized form), and coenzyme Q9 (reduced form)**

| Analytes | Precursor ion (*m/z*) | Fragment ion (*m/z*) | Polarity* | Collision Energy (eV) |
|---|---|---|---|---|
| Coenzyme Q10 (oxidized form) | 880.7 | 197.0 | $ESI^+$ | 20 |
| Coenzyme Q10 (reduced form) | 882.7 | 197.0 | $ESI^+$ | 30 |
| Coenzyme Q9 (oxidized form) | 812.6 | 197.0 | $ESI^+$ | 20 |
| Coenzyme Q9 (reduced form) | 814.6 | 197.0 | $ESI^+$ | 30 |

*ESI: Electrospray ionization (ESI) interface to generate protonated ions $[M+H]^+$

**Material information**

| REAGENT or SOURCES | SOURCE | IDENTIFIER |
|---|---|---|
| **Antibodies** | | |
| Anti-V5 Tag Monoclonal Antibody (mouse) | Invitrogen | Cat # R960-25 |
| Goat Anti-Mouse IgG (H + L)-HRP conjugate | Bio-Rad Laboratories | Cat # 1706516 |
| Anti-rabbit IgG, HRP-linked Antibody | Cell Signaling Technology | Cat # 7074S |
| Alexa Fluor 488 IgG mouse | Invitrogen | Cat # A11001 |
| Alexa Fluor 568 IgG mouse | Invitrogen | Cat # A11004 |
| Anti-RTN4IP1 | Atlas Antibodies | Cat # HPA036357 |
| Anti-TOM20 | ProteinTech | Cat # 11802-1-AP |
| p44/42 MAPK (Erk1/2) | Cell Signaling Technology | Cat # 9102 |
| Anti-8-OHdG | Santa Cruz Biotechnology | Cat # sc-66036 |
| **Chemicals, Peptides, and Recombinant Proteins** | | |
| Streptavidin-HRP | Thermo Fisher Scientific | Cat # 21126 |
| Streptavidin, Alexa Fluor 647 conjugate | Invitrogen | Cat # S21374 |
| Sodium 2,6-dichloroindophenolate hydrate (DCPIP) | Sigma-Aldrich | Cat # 119814 |
| NADPH | Sigma-Aldrich | Cat # 2646-71-1 |
| Biotin | Alfa Aesar | Cat # A14207 |
| $H_2O_2$ | Sigma-Aldrich | Cat # STBJ2658 |
| RIPA lysis buffer | ELPISBIO | Cat # EBA-1149 |
| Glutaraldehyde | Electron Microscopy Sciences | Cat # 16200 |
| Protease inhibitor cocktail | Invitrogen | Cat # 78438 |
| DAB | Sigma-Aldrich | Cat # D8001 |

| | | |
|---|---|---|
| Urea | Sigma-Aldrich | Cat # U5378 |
| Uranyl acetate | Electron Microscopy Sciences | Cat # 22400 |
| Embed-812 | Electron Microscopy Sciences | Cat # 14120 |
| Uranyless | Electron Microscopy Sciences | Cat # 22409 |
| Lead citrate | Electron Microscopy Sciences | Cat # 22410 |
| Lipofectamine 2000 | Life Technologies | Cat # 11668019 |
| 20X TBS | Thermo Fisher Scientific | Cat # 28358 |
| Acetone | Sigma-Aldrich | Cat # 650501 |
| Ammonium bicarbonate | Sigma-Aldrich | Cat # A6141 |
| Doxycycline | Sigma-Aldrich | Cat # D9891 |
| TPCK-Trypsin | Thermo Fisher Scientific | Cat # 20233 |
| Dithiothreitol | Sigma-Aldrich | Cat # 43819 |
| Iodoacetoamide | Sigma-Aldrich | Cat # I1149 |
| $CaCl_2$ | Alfa Aesar | Cat # 12312 |
| Trifluoroacetic acid | Sigma-Aldrich | Cat # T6508-10AMP |
| Formic acid | Thermo Fisher Scientific | Cat # 28905 |
| Acetonitrile | Sigma-Aldrich | Cat # 900667 |
| Coenzyme Q10 (oxidized form, >98%) | Sigma-Aldrich | Cat # C9538 |
| Coenzyme Q9 (oxidized form, >98%) | ChemScene | Cat # CS-6359 |
| Coenzyme Q10 (reduced form, >95%) | Biosynth Carbosynth | Cat # FU28634 |
| **Critical Commercial Assays** | | |
| TMRE | Sigma-Aldrich | Cat #115532-52-0 |
| Seahorse XF Cell Mito Stress Test Kit | Agilent | Cat #103015-100 |
| Seahorse XF DMEM medium | Agilent | Cat #103575-100 |
| Seahorse XF 1.0 M glucose solution | Agilent | Cat #103577-100 |
| Seahorse XF 100mM pyruvate solution | Agilent | Cat #103578-100 |
| Seahorse XF 200mM glutamine solution | Agilent | Cat #103579-100 |
| **Oligonucleotides** | | |

| | | |
|---|---|---|
| Small interfering RNA (siRNA) for *Rtn4ip1* | Bioneer | Cat # 170728-1 |
| Primers for genotyping (5’-3’) F:GGTACCATGCTGGCCACCCGCGTGTTC R:GCGGCCGCTCATTAGGCATCAGCAAACCCAAGCTCGGAA | This paper | NA |
| **Recombinant DNA** (See detail information in **Table S7** of Supplemental Information) | | |
| MTS-V5-APEX2_pCDNA5 | Lee et al., 2017 | NA |
| RTN4IP1-V5-APEX2_pCDNA5 | This paper | NA |
| MTS (from RTN4IP1)-V5-APEX2_pCDNA5 | This paper | NA |
| RTN4IP1(Δ1-32aa)-V5-APEX2_pCDNA5 | This paper | NA |
| His6_MBP_TEV_ RTN4IP1(Δ1-32aa)_pET21a | This paper | NA |
| RTN4IP1-TurboID_pCDNA5 | This paper | NA |
| MTS-V5-TurboID_pCDNA5 | This paper | NA |
| lentiCRISPRv2 | Addgene | Cat #52961 |
| **Software and Algorithms** | | |
| Morpheus | Broad Institute | https://software.broadinstitute.org/morpheus/ |
| STRING | CPR, EMBL, SIB, KU, and UZH | https://string-db.org/ |
| MitoFates | AIST | http://mitf.cbrc.jp/MitoFates/cgi-bin/top.cgi |
| Image J | NIH | https://imagej.nih.gov/ij/ |
| Pymol | Schrodinger | https://pymol.org/2/ |
| Normalizer | Lund University, Medicon Village 406, 223 81, Lund, Sweden | http://normalyzer.immunoprot.lth.se/ |
| Perseus | Max-Planck-Institute of Biochemistry | https://maxquant.net/perseus/ |
